# Conoid extrusion serves as gatekeeper for entry of glideosome components into the pellicular space to control motility and invasion in Apicomplexa

**DOI:** 10.1101/2022.06.06.494926

**Authors:** Nicolas Dos Santos Pacheco, Lorenzo Brusini, Romuald Haase, Nicolò Tosetti, Bohumil Maco, Mathieu Brochet, Oscar Vadas, Dominique Soldati-Favre

**Author notes:** Equal contribution (Lorenzo Brusini, Romuald Haase).

## Abstract

Members of the Apicomplexa are defined by apical cytoskeletal structures and secretory organelles, tailored for motility and invasion. Gliding is powered by actomyosin-dependent rearward translocation of apically secreted transmembrane adhesins. In *Toxoplasma gondii*, the conoid, composed of a cone of spiraling tubulin fibers and apposed preconoidal rings (PCRs), is an enigmatic, dynamic organelle of undefined function. Here we mapped five new components of the PCRs and deduce that the structure serves as a pivotal hub for actin polymerization and glideosome assembly. F-actin produced by Formin1 on the PCRs is used by Myosin H to generate the force for conoid extrusion. A set of B-box-type zinc finger domain containing proteins conserved in Apicomplexa is indispensable for PCRs formation, conoid extrusion and motility in *Toxoplasma* and *Plasmodium*. Conoid dynamics directs the flux of F-actin to the pellicular space, acting as dynamic gatekeeper to tightly control parasite motility during invasion and egress.

## INTRODUCTION

The phylum of Apicomplexa includes a large group of eukaryotic parasites of considerable medical and veterinary importance ^1^. Human parasites *Toxoplasma gondii, Plasmodium falciparum* and *Cryptosporidium* species are responsible for toxoplasmosis, malaria, and cryptosporidiosis, respectively ^2-5^. These parasites depend on host cell invasion for survival, replication, and dissemination. Invasion and egress from infected cells as well as crossing biological barriers rely on gliding motility ^6^, itself dependent on a series of conserved cytoskeletal components and secretory organelles collectively known as the apical complex ^7, 8^.

As members of Alveolata, apicomplexans possess a pellicle composed of the plasma membrane (PM) and inner membrane complex (IMC) formed of set of armor-plate like sacs also named alveoli ^9^. The IMC is supported by the subpellicular network (SPN) composed of intermediate-like filaments ^10^ and an array of subpellicular microtubules (SPMTs) ^11^ emerging from an apical polar ring (APR), a circular-shaped microtubule-organizing center (MTOC) ^12, 13^. Host cell invasion involves secretion of specialized apical organelles termed micronemes and rhoptries ^14, 15^. The transmembrane adhesin Apical Membrane Antigen 1 (AMA1), secreted by micronemes to the surface of the parasite, interacts with a complex of discharged rhoptry neck proteins (RON complex) serving as receptors at the surface of the host cell ^16^. The glideosome-associated connector (GAC) physically links adhesins to actin filaments (F-actin) ^17^ generated by Formin 1 (FRM1), an essential actin nucleator localizing at the apical tip of Apicomplexa ^18-20^. Translocation of adhesins to the posterior pole is driven by the apico-basal flux of F-actin to produces forward motion, allowing penetration of the parasite inside the forming parasitophorous vacuole (PV) ^6, 21^. The flux of F-actin is generated by the concerted action of myosin H (MyoH) at the conoid, and myosin A (MyoA), anchored between the IMC and parasite PM ^18, 22, 23^. An apical lysine methyltransferase (AKMT) critically participates in the regulation of F-actin flux and the apical recruitment of GAC by as yet unknown mechanisms ^17, 18, 24^. These key invasion factors concentrate at the apical tip, however their precise arrangement to orchestrate glideosome assembly remains unknown.

Positioned in the vicinity of the APR, the conoid is a dynamic organelle composed of curved tubulin fibers forming an open cone, topped by preconoidal rings (PCRs) and traversed by two intraconoidal microtubules (ICMTs) ^8^. Recent studies identified multiple conoid proteins in *Plasmodium* species and demonstrated the existence of a reduced conoid lacking ICMTs ^25-28^. In *Toxoplasma*, the conoid extrudes and retracts through the APR in motile parasites, but how and why these movements take place is unclear ^29-31^ and whether conoid extrusion is required for microneme secretion ^32^ or rhoptry discharge is not known ^33, 34^.

Here, we used Ultrastructure Expansion Microscopy (U-ExM) to unambiguously assign the suborganellar localization of key known conoid proteins and reliably assess conoid extrusion in *T. gondii*. The presence of FRM1, AKMT and GAC at the PCRs establishes that this structure serves as platform for actin polymerization and glideosome assembly. The identification, biochemical, structural, and functional characterization of two novel components of the PCRs also demonstrate a pivotal role in gliding motility. Pcr4 and Pcr5 assemble in stable heterodimers *in vitro* and form the core of PCRs *in vivo*, while Pcr6 is required tom tether PCRs to the conoid. Pcr4 and Pcr5 are conserved in Apicomplexa and are also required for *Plasmodium berghei* ookinete motility. Finally, MyoH powers conoid extrusion, a process that serves as gatekeeper for the entry of F-actin in the pellicular space ensuring a tight regulation of gliding motility during invasion and egress.

## RESULTS

### The preconoidal rings serve as a hub for actin polymerization and glideosome assembly

Several key proteins implicated in gliding motility and invasion have previously been shown to localize to the apical tip of the parasite by indirect immunofluorescence assay (IFA) ^17, 18, 23, 24, 35^. In contrast to conventional microscopy, tubulin staining visualized by U-ExM unambiguously resolved the conoid complex with tubulin fibers and ICMTs distinct from subpellicular microtubules ^36^. SAS6L, DCX and CPH1 were confirmed to localize to the cone ^37-39^, as well as MyoH for which the cone localization was already suggested previously ^23^ (Fig.1a and Extended Fig.1a). In contrast, FRM1, GAC and the apical signal of Centrin2 (CEN2) formed a ring above the conoid, reminiscent of the PCRs (Fig.1a). AKMT, localizing apically exclusively in intracellular parasites, was found both at the PCRs and the cone in intracellular parasites (Extended Fig.1b) and non-activated extracellular parasites resuspended in a buffer mimicking intracellular conditions ^24^ (Fig.1a). Consistent with AKMT localization, the PCRs were heavily stained by an anti-methylated lysine antibody ^17^ (Fig.1a).

**Figure 1.**
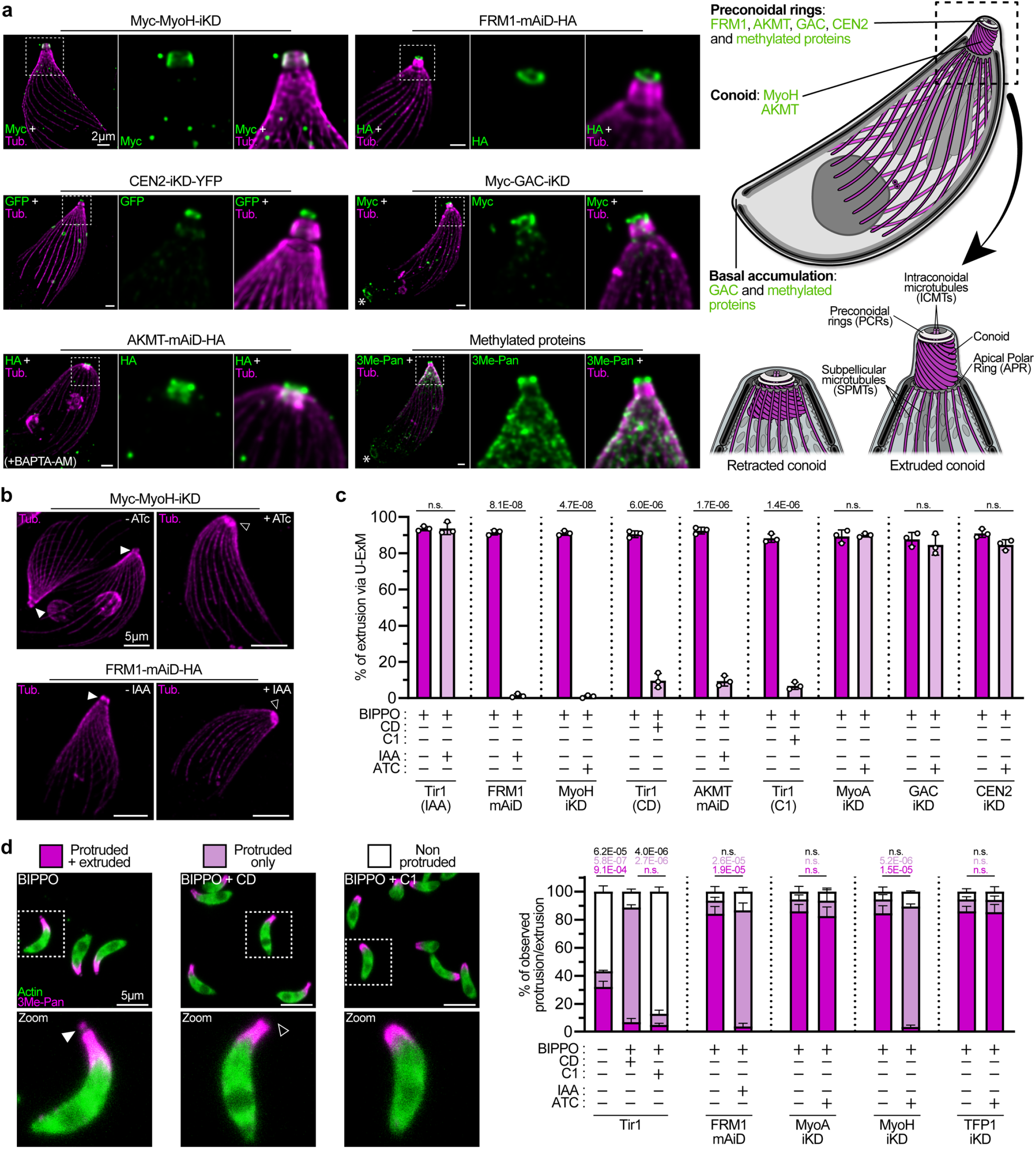
Conoid extrusion is an actomyosin dependent process powered by MyoH and fueled by FRM1. **(a)**Localisation of apical protein via U-ExM. Asterisks indicate the basal pole of parasites. Tub = tubulin. Scale bars = 2 µm. **(b)** Representative pictures of conoid extrusion assessment via U-ExM. White arrowhead = extruded conoid. Black arrowhead = retracted conoid. Scale bars = 5 µm. **(c)** Quantification of conoid extrusion via U-ExM as displayed in Fig.1b. For practicality, -mAiD stands for - mAiD-HA strains and -iKD stands for Tet-fused genes of interest. For each condition, 200 parasites were counted in three biologically independent experiments (n=3). Unpaired t-test where non-significant (n.s.) if P>1E-02. **(d)** Protrusion/extrusion assay by regular IFA. White arrowhead = extruded conoid. Black arrowhead = retracted conoid. Scale bars = 5 µm. On the right, quantification of conoid extrusion and parasite protrusion. For each condition, 200 parasites were counted in three biologically independent experiments (n=3). Unpaired t-test where non-significant (n.s.) if P>5E-02.

### Conoid extrusion is powered by myosin H and F-actin generated by Formin 1

Previously, visualizing conoid movement has been limited to a change in *Toxoplasma* shape as observed by phase contrast ^29, 31^. Parasites displaying an elongated apical shape were considered as “extruded” despite no clear visualization of the conoid. Upon stimulation of extracellular parasites with a phosphodiesterase inhibitor (BIPPO) ^40^, U-ExM unambiguously defined conoid extrusion when the cone is seen above the APR and SPMTs basket (Fig.1b and Extended Fig.1c). Depletion of either FRM1 or MyoH completely abrogated conoid extrusion (Fig.1b,c and Extended Fig.1c), confirming this is an actomyosin-dependent process as previously anticipated by treatment with cytochalasin D (CD), an inhibitor of actin polymerization ^29, 31^. Treatment with Compound 1, an inhibitor of the cyclic GMP-dependent protein kinase, PKG, which is essential for microneme secretion, motility, invasion, and egress, blocked conoid extrusion (Fig.1c). Depletion of AKMT, previously shown to interfere with the apico-basal flux of F-actin ^18^ also blocked conoid extrusion (Fig.1c). In contrast, depletion of MyoA or GAC that are crucial for parasite motility but dispensable for initiation of F-actin flux, did not alter conoid extrusion (Fig.1c). Depletion of CEN2, a protein localized to the preconoidal rings and implicated in microneme secretion ^35, 41, 42^ did not affect conoid extrusion either (Fig.1c). To discriminate between parasite elongation and conoid extrusion and to provide a rationale for previous erroneous interpretation, we took advantage of the dual localization of anti-methylated lysine antibody (3Me-Pan) ^17^. Conoid extrusion can be assessed via the staining of an apical dot reminiscent of the PCRs, while the apical cap staining assessed the protruded or round shape of the parasites (Fig.1d). While CD blocked conoid extrusion only, C1 affected both conoid extrusion and parasite elongation. As observed by U-ExM, depletion of FRM1 or MyoH abrogated conoid extrusion only, while depletion of MyoA affected neither conoid extrusion nor parasite elongation (Fig.1d). In addition, using TFP1-depleted parasites lacking mature micronemes ^43^, extrusion and elongation were not affected, indicating that both processes are not linked to microneme secretion (Fig.1d).

### Identification of key components of the preconoidal rings

Hyperplexed localization of organelle proteins by isotope tagging (hyperLOPIT) has previously assigned *T. gondii* proteins to sub-cellular compartments ^44^, identifying two clusters enriched in proteins localized at the apical complex. Combining this data with cell cycle dependent transcription profiles and fitness conferring scores ^45^ we further identified five proteins newly localized to the PCRs called Pcr (<u>P</u>re<u>c</u>onoidal rings) (Fig.2a). Pcr5 belongs to a family of proteins that possess two B-Box zinc finger domains ^46^. The genome of *T. gondii* encodes four other proteins containing B-Box domains including two found at the PCRs, namely Pcr4 and Pcr7. Upon C-terminal fusion with a mini-auxin degron (mAiD) and a triple HAtag at their respective endogenous locus, the proteins showed apical dots by regular IFAs, except for Pcr7 which displayed an apical localization only in nascent daughter cells (Extended Fig.2a). When assessed by U-ExM, Pcr4, Pcr5 and Pcr6 localized to the PCRs (Fig.2a and Extended Fig.2b). In contrast, Pcr1 that was already localized at the apical pole ^26, 47^ displayed a unique signal, forming a “half-ring” at the level of PCRs, while Pcr7 localized exclusively to the PCRs of nascent daughter cells.

**Figure 2.**
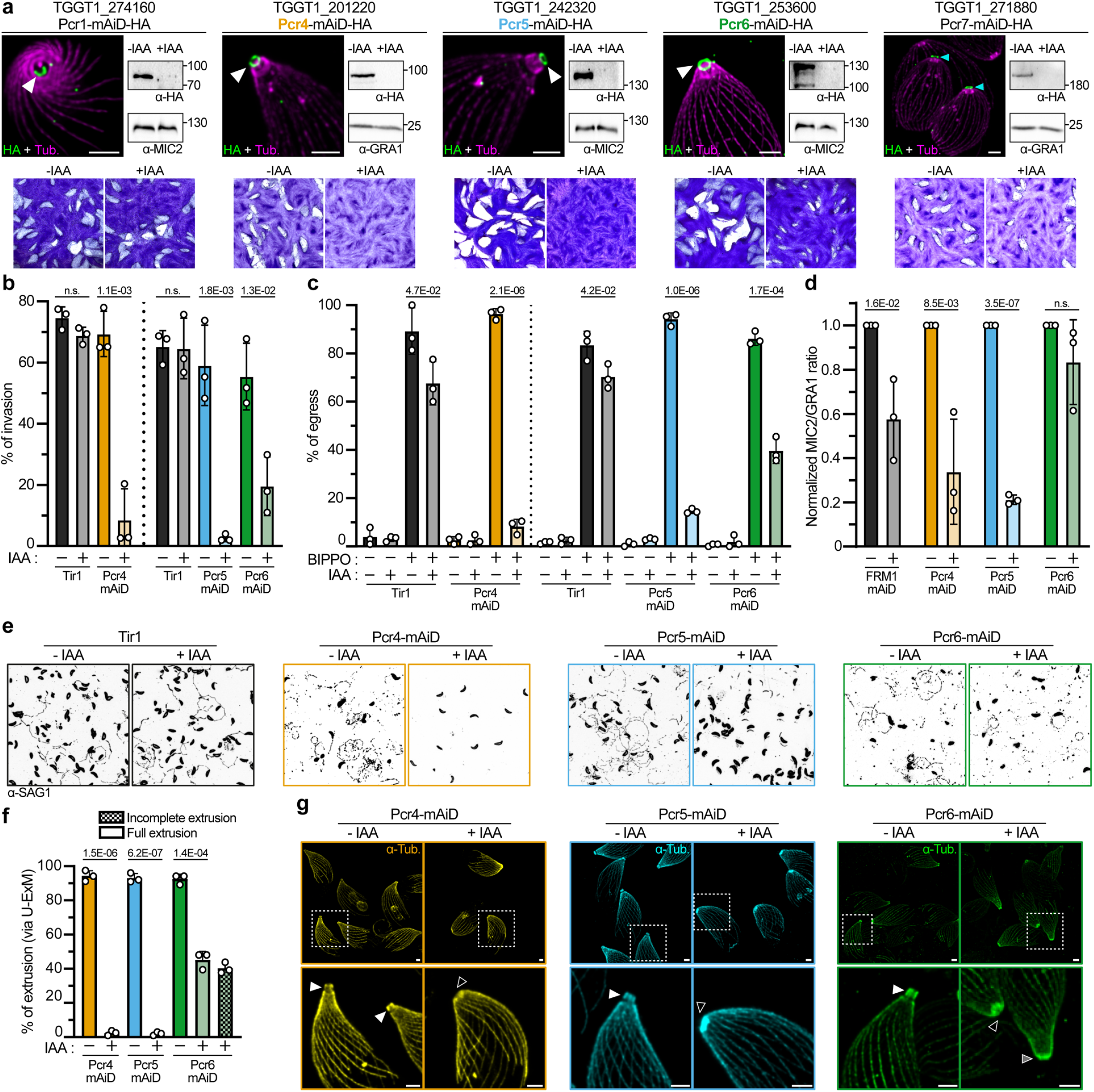
Five new preconoidal ring proteins. **(a)** Localization by U-ExM, regulation by Western Blot and plaque assay of the five proteins fused with a mAiD-HA cassette. White arrowhead = apical signal. Cyan arrowhead = apical signal in forming daughter cells. Scale bars = 2 µm. **(b)** Red/green invasion assay. **(c)** Induced egress assay. **(d)** Quantification of microneme secretion as assessed by band densitometry. **(e)** Trail deposition assay (gliding assay). Parasites and trails are labeled with SAG1 antibody. **(f)** Conoid extrusion assessed via U-ExM. **(g)** Representative pictures of panel (f). White arrowhead = extruded conoid. Black arrowhead = retracted conoid. Grey arrowhead = half-extruded conoid. Scale bars = 2 µm. For all statistical analysis, an unpaired t-test where non-significant (n.s.) if P>5E-02 was used. All data are presented as mean ± SD (n=3).

Depletion of protein showed that Pcr4 and Pcr5 are individually essential for parasite survival, forming no plaque of lysis 7 days after inoculation on monolayers of human foreskin fibroblasts (HFF) in presence of auxin (IAA) (Fig.2a). Conditional depletion of Pcr6 exhibited an intermediate phenotype with the formation of plaques 60% smaller while depletion of either Pcr1 or Pcr7 did not show any reduction in plaque size, indicating that these proteins are possibly dispensable (Fig.2a and Extended Fig.2c).

Pcr4 and Pcr5 are homologous. Homologs for Pcr4 and Pcr5 were detected across Apicomplexa and Dinoflagellates, with Pcr4 homologs further distributed in ciliates, indicating these genes duplicated before the last common ancestor of Myzozoa (Extended Fig.3a,b). In contrast, Pcr6 was not clearly detected outside Apicomplexa.

### Pcr4, Pcr5 and Pcr6 contribute to egress, motility and invasion

In contrast to previously reported mAiD tagged proteins that are targeted for degradation within less than one hour ^48^, Pcr4 and Pcr5 required 8 hours incubation in presence of IAA to be depleted beyond detection (Extended Fig.4a) and both proteins are stable in non-dividing extracellular parasites incubated with IAA (Extended Fig.4b). Taken together, these results suggest that Pcr4 and Pcr5 are irreversibly assembled in the PCRs and their turnover only occurs during parasite division. In good agreement, short auxin treatment (< 6 hours) only affected forming daughter cells PCRs proteins, while long auxin treatment (> 6 hours) also affected mature cells (Extended Fig.4c). IFA also established that Pcr4, Pcr5 and FRM1 were detectable before ISP1, one of the earliest known markers of daughter cell formation ^49^. In contrast, Pcr6 was never observed before the appearance of ISP1, highlighting the temporal hierarchy in the incorporation of components at the PCRs. Depletion of Pcr4 and Pcr5 led to severe defects in invasion and in egress at a rate of less than 10 % of the parental cell lines, while depletion of Pcr6 led to an intermediate phenotype of 20% in invasion and 40% in egress (Fig.2b,c). To determine if the impairments in invasion and egress could be attributed to a defect in microneme secretion, an assay based on quantification of excreted secreted antigens (ESA) by western blot was performed ^50^. While Pcr4 and Pcr5 mutants exhibited significant decrease in microneme secretion (Fig.2d and Extended Fig.4d), no full abrogation of exocytosis was observed. This contrasts with more severely impacted mutants interfering with the cGMP signaling cascade ^51^, microneme morphogenesis ^43^ or loss of the conoid complex ^48, 52^. Remarkably, FRM1 depleted parasites only showed a decrease of 40 % in microneme secretion, despite a drastic impairment in conoid extrusion. For comparison, depletion of Pcr4 and Pcr5 led to decrease of 70-80% in secretion.

A standard trail deposition assay showed that Pcr4-2 are required for gliding motility, while Pcr6-lacking parasites are only able to form circular trails (Fig.2e). U-ExM further revealed the conoid failed to extrude in Pcr4 or Pcr5 depleted parasites (Fig.2f,g). In Pcr6 depleted parasites, conoid extrusion was reduced to 40% with the majority of parasites exhibiting neither fully extruded nor fully retracted conoids. Taken together, these results indicate that Pcr4 and Pcr5 are essential for conoid extrusion, an event not absolutely required for microneme exocytosis.

### Pcr4, Pcr5 and Pcr6 are involved in the assembly of the PCRs

To further determine the hierarchy of PCR components, we looked at the recruitment and co-dependency of Pcr4 relative to Pcr5, in cells depleted for either protein. In both cases, when one protein was downregulated, the other one could not be visualized by IFA anymore (Fig.3a). Similarly, Pcr6 and Pcr1 were lost in the absence of Pcr5 (Fig.3b,c), whilst depletion of Pcr6 did not impact on Pcr5 (Extended Fig.5a). Western blot analysis revealed a decrease in levels of Pcr6 and Pcr1 upon Pcr5 depletion. Importantly, levels and localization of FRM1 were completely abolished upon Pcr5 depletion (Fig.3d), whereas Pcr4 and Pcr5 were not affected by depletion of FRM1 (Extended Fig.5b). In the case of CEN2, only the apical localization was lost upon Pcr5 depletion, whereas staining at peripheral annuli, centrioles and the basal complex was retained ^41^ and levels apparently not affected (Fig.3e). Markers of the APR, namely RNG1 and KinA, or the cone of the conoid, like Cam2, were unaffected upon depletion of Pcr5 (Extended Fig.5c). Finally, depletion of Pcr5 led to the loss of AKMT and GAC at the PCRs but did not clearly affect signal at the cone (Extended Fig.5d,e). Negative-stain transmission electron microscopy (TEM) further characterized the effect of PCRs protein depletion. In absence of either Pcr4 or Pcr5, while very few parasites displayed an extruded conoid, the ICMTs and APR were still present but the PCRs were visibly lost (Fig.3f and Extended Fig.5f). In contrast, depletion of Pcr6 led to PCRs detaching from the cone (Fig.3f and Extended Fig.5g), whereas absence of FRM1 had no clear effect on the ultrastructure of the apical complex (Extended Fig.5h). Altogether, Pcr4 and Pcr5 are essential for the formation, and maintenance of the PCRs that serve as anchor for FRM1, CEN2, GAC and AKMT, the key players in actin polymerization, microneme secretion and glideosome assembly, respectively.

**Figure 3.**
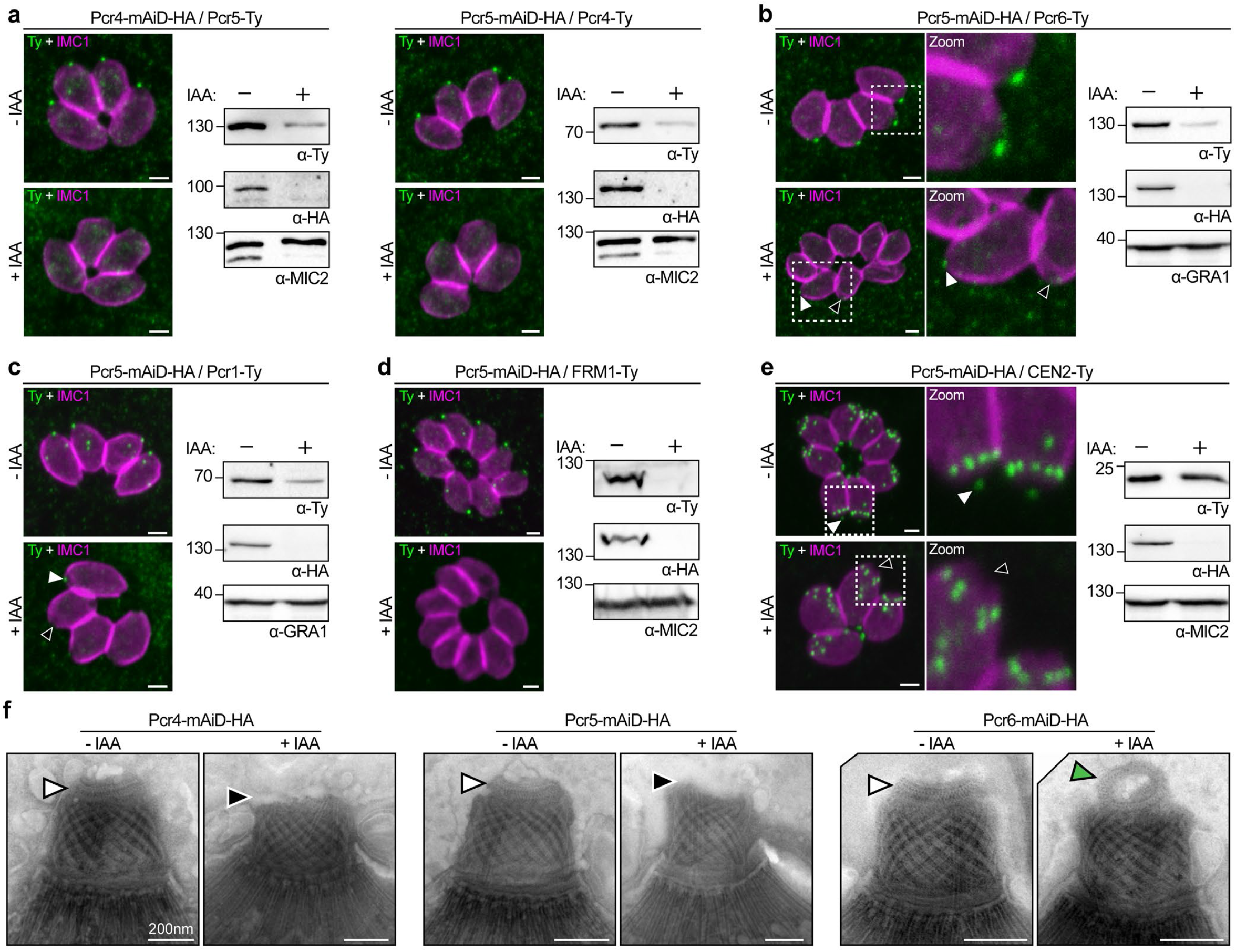
Depletion of Pcr4 or Pcr5 causes the loss of the preconoidal rings. **(a)** Effect of Pcr4 depletion on Pcr5 and conversely. **(b)** Effect of Pcr5 depletion on Pcr6. White arrowhead = presence of apical signal. Black arrowhead = no apical signal. **(c)** Effect of Pcr5 depletion on Pcr1. White arrowhead = presence of apical signal. Black arrowhead = no apical signal. **(d)** Effect of Pcr5 depletion on FRM1. **(e)** Effect of Pcr5 depletion on Centrin2. White arrowhead = presence of apical signal. Black arrowhead = no apical signal. All scale bars = 2 µm. **(f)** Effect of Pcr4, Pcr5 or Pcr6 depletion on preconoidal rings as seen by electron microscopy. White arrowhead = presence of preconoidal rings. Black arrowhead = absence of preconoidal rings. Green arrowhead = detached preconoidal rings. Scale bars = 200 nm.

### Pcr4 and Pcr5 form a stable heterodimer *in vitro* and the B-Box domains of Pcr5 participate in preconoidal rings assembly *in vivo*

B-Box zinc-finger domains are characterized by eight conserved cysteines and/or histidine residues that chelate two zinc ions and are typically implicated in nucleic acid-protein or protein-protein interactions ^46, 53^. Pcr4 and Pcr5 likely originate from gene duplication and each possess two B-Box domains (Extended Fig.6a). Given their sequence similarity, individual essentialities and comparable phenotypes upon depletion, along with the presence of coiled-coil domains, it is possible these proteins also interact biochemically. To determine their biochemical properties, Pcr4 and Pcr5 were expressed both individually and together in baculovirus-infected insect cells. Whilst purified Pcr5 formed large oligomers when expressed alone, Pcr4 was only soluble when co-expressed with Pcr5, yielding a soluble hetero-complex (Fig.4a-c and Extended Fig.6b-d). Analysis of the Pcr4-Pcr5 complex by pull-down, mass photometry or size-exclusion chromatography coupled with multi-angle light scattering (SEC-MALS) indicated both proteins formed a stable and homogeneous heterodimer in solution (Extended Fig.6e,f). AlphaFold ^54^ further predicted Pcr4 and Pcr5 to adopt similar conformations together forming an antiparallel heterodimer, joined by a central coiled-coil region that is flanked by the tandem B-Box domains of each protein (Fig.4d,e). Both proteins encode an N-terminal antiparallel beta-strand domain that packs against the central coil-coil and Pcr5 harbors an additional C-terminal alpha-helical domain that contacts the Pcr4 beta-strand domain. Consistent with structure prediction, hydrogen/deuterium exchange coupled to mass spectrometry (HDX-MS) confirmed that residues within centrally predicted coiled-coil domains of Pcr4 and Pcr5 have low H/D exchange rate ^55^ (Fig.4d,f and Supplementary Table 1). Beta-strand domains of both proteins have intermediate H/D exchange rates and the C-terminal alpha-helical region of Pcr5 shows both low and high H/D exchange rates. Overall, the HDX-MS experimental results compellingly support the AlphaFold model, with regions having high dynamics corresponding to regions with poor structural prediction score (pLDDT score) (Fig.4d).

**Figure 4.**
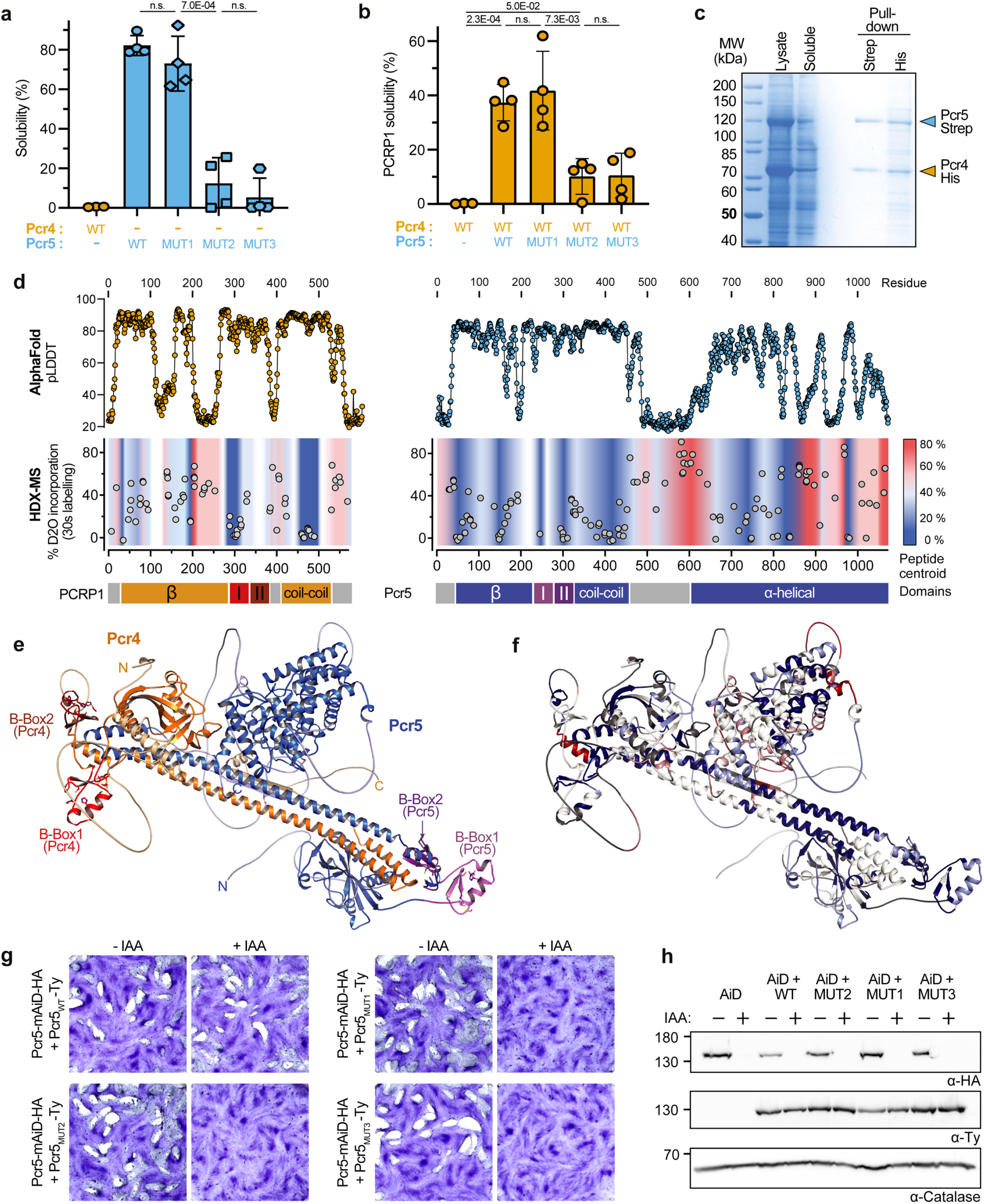
Pcr4 and Pcr5 form a complex in vitro. **(a)** Solubility of recombinant isolated Pcr4 and the four mutated versions of recombinant isolated Pcr4 and the four mutated versions of Pcr5. Solubility was assessed by band densitometry comparing cell lysates before and after centrifugation on Coomassie-stained SDS-PAGE gels. **(b)** Solubility assay of Pcr4 in presence of the four versions of Pcr5. **(c)** Pull down assay of Pcr4-His + Pcr5-Strep when both proteins were co-expressed. **(d)** Graphical representation of AlphaFold confidence score (pLDDT), HDX-MS data presenting the exchange dynamic (D_2_0 incorporation), and the protein architecture and domains. **(e)** AlphaFold prediction of the Pcr4-Pcr5 complex 3D structure. **(f)** HDX-MS data (panel d) plotted on the AlphaFold prediction of the → Pcr4-Pcr5 complex. **(g)** Plaque assay of the four complemented strains. **(h)** Western blot analysis of the four complemented strains. HA = endogenous copy. Ty = second copy. Catalase = loading control. For all statistical analysis, an unpaired t-test where non-significant (n.s.) if P>5E-02 was used. All data are presented as mean ± SD (n=3-4).

To assess the importance of Pcr5 B-Box domains towards the formation of the Pcr4-Pcr5 complex *in vitro*, zinc-chelating residues were mutated to alanine (Extended Fig.6a). While Pcr5_MUT1_ was soluble, Pcr5_MUT2_ and Pcr5_MUT3_ (mutations in both B-Box domains) were insoluble, suggesting that an intact second B-Box domain is required for proper protein conformation (Fig.4a,b). However, the B-Box domains of Pcr5 are not implicated in the Pcr4-Pcr5 complex formation *in vitro* as both proteins could be isolated by pull-down of wildtype or mutated versions of Pcr5 (Extended Fig.6g). HDX-MS further showed that the Pcr4Pcr5_MUT1_ recombinant complex had an almost identical conformation compared to non-mutated complex, with only the B-Box 1 region of Pcr5 showing different dynamics in the mutant (Extended Fig.6h).

To further investigate the possibility of loss-of-function mutations within the B-Box domains of Pcr5 *in vivo*, C-terminally tagged Pcr5 ORFs were ectopically expressed in the presence and absence of endogenous Pcr5. While parasites expressing a second copy of Pcr5_WT_ completely rescued the effect of the endogenous Pcr5 depletion, mutated second copies were unable to (Fig.4g,h). IFA confirmed Pcr5_WT_ localized both to the cytosol and the apical tip, whereas mutant copies did not localize to the apical tip (Extended Fig.6i). Of relevance, all second copies of Pcr5 were capable of retaining endogenous Pcr5 in the cytosol (Extended Fig.6i), without changing protein levels (Fig.4h). Taken together the B-Box domains fulfill an important role is assembly of the PCRs that is unrelated to Pcr4-Pcr5 conformation.

### Pcr4 and Pcr5 have a conserved role in *Plasmodium berghei* ookinetes

Until recently, the conservation of the conoid in the related apicomplexan models and malaria parasites, i.e. *Plasmodium*, was limited to EM images of the apical end of invasive zoites ^56, 57^. The presence of a reduced conoid in ookinete stages of *P. berghei* was only shown recently ^25, 26, 28^. To investigate if Pcr4 and Pcr5 also localize to the PCRs in malaria parasites, both proteins were localized at multiple invasive stages throughout the *Plasmodium* lifecycle by integration of the coding sequence of mNeonGreen fused to triple-HA tag at the endogenous locus of each gene. Live-fluorescence imaging revealed both PbPcr4 and PbPcr5 first accumulated as punctae in late asexual-blood stages during trophozoite development (Fig.5a), which were maintained following progression to schizont stages and formation of daughter cell bodies, with one clear punctum present at the surface of each fully formed and extracellular merozoite. Consistent with the notion that all invasive *Plasmodium* forms possess PCRs, a similar signal was present at the apical end of ookinetes. U-ExM further and unambiguously determined PbPcr4 and PbPcr5 localize to a small ring above the ookinetes conoid (Fig.5b). As PCRs serve as anchor for several key glideosome components in *Toxoplasma*, PbGAC-GFP ^17^ localized similarly at PCRs of ookinetes (Fig.5b).

**Figure 5.**
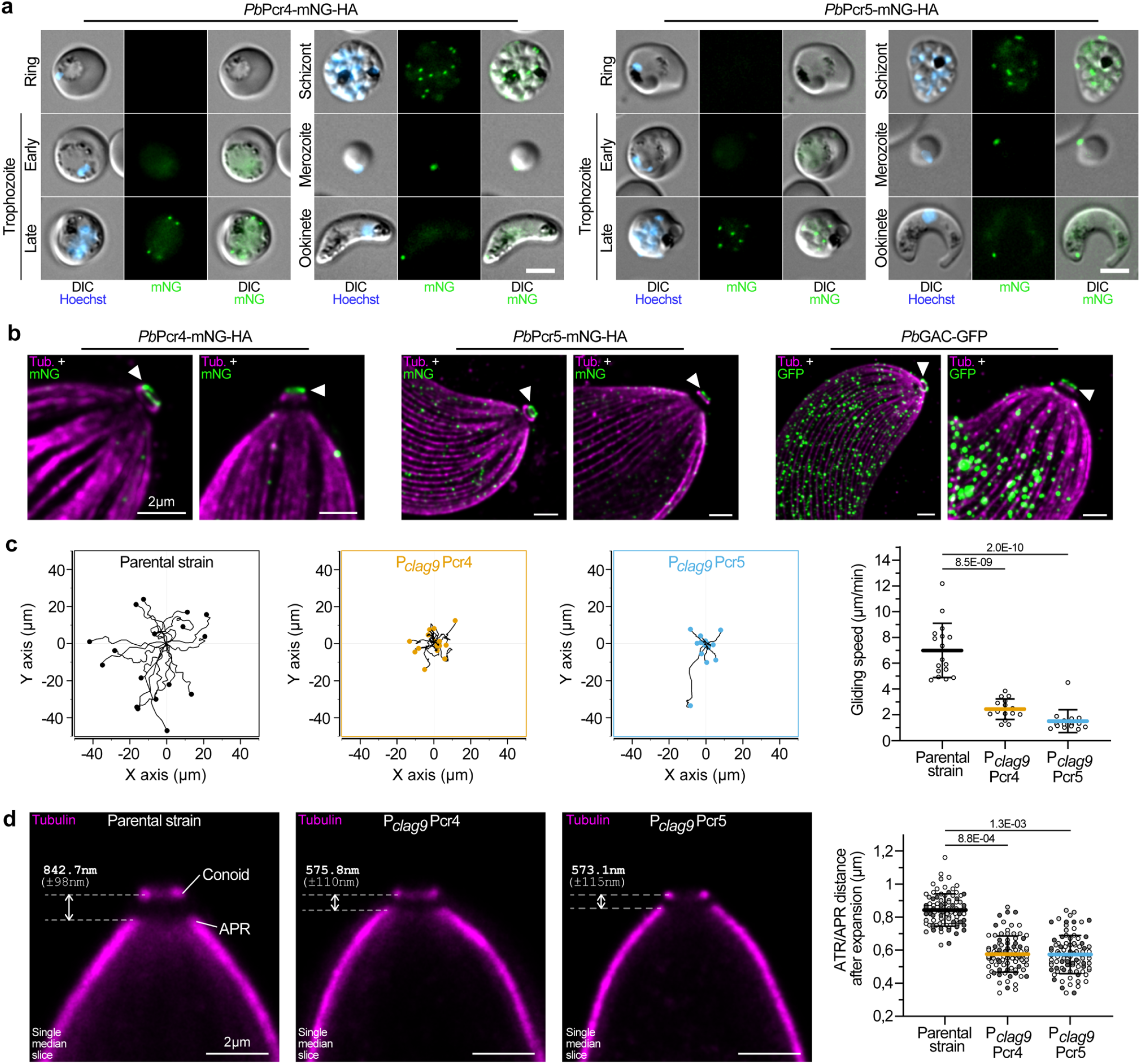
Pcr4 and Pcr5 localization and phenotypes are recapitulated in *P. berghei*. **(a)** Localization of *Pb*Pcr4 and *Pb*Pcr5 by live-microscopy. Scale bars for ookinete = 5µm. **(b)** Localization of *Pb*Pcr4, *Pb*Pcr5, and *Pb*GAC in *P. berghei* ookinetes by U-ExM. White arrowhead = preconoidal ring staining. Scale bars = 2 µm. **(c)** Ookinete gliding motility assessed by the tracking of individual cells by live microscopy (all individual cells were tracked over n=3 independent biological replicates). On the right panel, gliding speed from individual ookinetes was also quantified. **(d)** Representative U-ExM pictures of ookinete apical pole. Right panel show the measurement of the conoid-APR distance after expansion (gels of identical expansion ratio were used). (n=3 independent biological replicates represented by 3 shades of grey). Scale bars = 2 µm. For all statistical analysis, an unpaired t-test where non-significant (n.s.) if P>5E-02 was used. All data are presented as mean ± SD (n=3).

Genome-wide screenings have indicated Pcr4 and Pcr5 are essential for blood-stage proliferation in human and rodent malaria parasites ^58, 59^. To study the role of PbPcr4 and PbPcr5 during sexual development and mosquito stages, both genes were placed under the control of the *clag9* promoter (P_*clag9*_Pcr4 and P_*clag9*_Pcr5) ^60^, to maintain relative levels of both proteins in asexual blood-stages but reducing expression in sexual stages (Extended Fig.7a). Whilst P_*clag9*_Pcr4 and P_*clag9*_Pcr5 clones readily formed microgametes and converted to banana-shaped ookinete forms compared to parental cell lines (Extended Fig.7b,c), ookinetes appeared less motile. Time-lapse movies of ookinetes in thin layers of Matrigel (Fig.5c and Supplementary Movies 1-3), revealed down-regulation of PbPcr4 or PbPcr5 resulted in a decrease in average gliding speed from 6.99 ± 2.11 µm/min in wild type ookinetes to 2.43 ± 0.80 µm/min and 1.50 ± 0.88 µm/min in P_*clag9*_Pcr4 and P_*clag9*_Pcr5 lines, respectively. Furthermore, tubulin staining in expanded cells revealed the distance of the conoid relative to the APR was markedly reduced in P_*clag9*_Pcr4 and P_*clag9*_Pcr5 lines compared to parental cells (576 and 573 nm compared to 843 nm, respectively) (Fig.5d). Taken together, the PCRs composition and function appear conserved across the phylum.

### Conoid extrusion gates F-actin to the pellicular space to regulate motility

Our data has led us to postulate a model to explain the role of conoid extrusion and retraction in controlling parasite motility and invasion (Fig.6a). In this model, extrusion of the conoid above the APR is a prerequisite to gate the flow of F-actin inside the pellicular space upon the action of MyoH. At the level of the APR, MyoA takes the relay to sustain the flux of F-actin toward the posterior pole of the parasite ^17, 23^. In combination with GAC and microneme secretion, the flux of F-actin results in productive motility.

**Figure 6.**
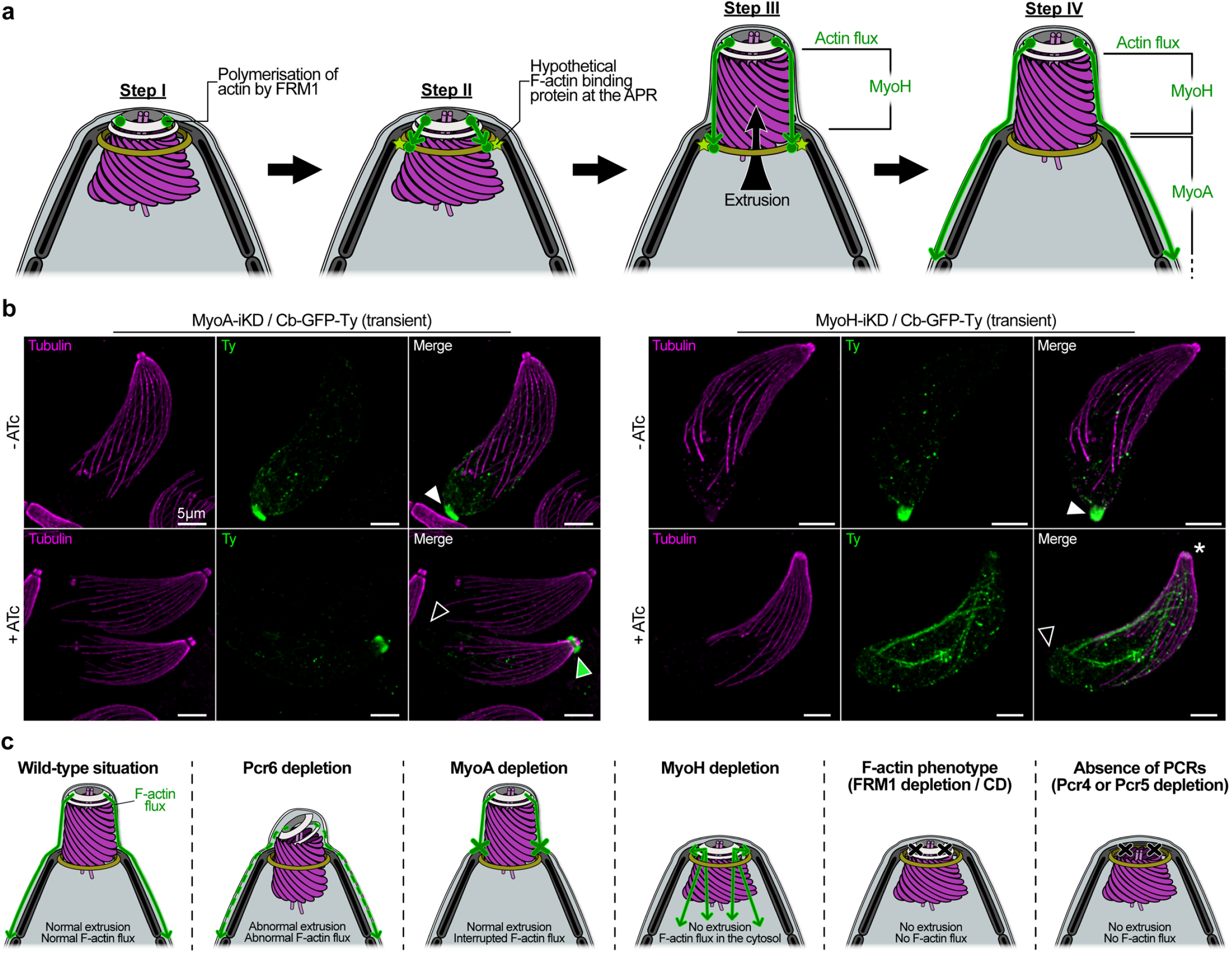
The preconoidal rings serve as a gatekeeper for glideosome entry in the pellicular space. **(a)** Hypothetical model for the conoid extrusion process. First, actin is polymerized by FRM1 at the level of the PCR. Then, the F-actin is translocated by MyoH towards the APR, where an hypothetical F-actin binding protein anchor the filaments to the APR. Once anchored, the action of MyoH will push the conoid forward to complete extrusion. Finally, F-actin is translocated first by MyoH at the conoid level, and then by MyoA at the pellicle to induce motility. **(b)** U-ExM on MyoA and MyoH mutants expressing transiently F-actin binding chromobodies (Cb-GFP-Ty). White arrowhead = F-actin basal accumulation. Black arrowhead = absence of F-actin basal accumulation. Green arrowhead = F-actin accumulation at the MyoH-MyoA relay. Asterisk = absence of conoid extrusion. Scale bars = 5µm. **(c)** Summary of the different phenotype observed in this study.

To challenge this model, we applied U-ExM on parasites transiently expressing F-actin-recognizing chromobodies fused to a GFP-Ty-tag (Cb-GFP-Ty) in cells depleted for MyoH or MyoA. In presence of both myosins, F-actin systematically accumulated at the basal pole of the parasites (Fig.6b,c) as previously reported ^18^. Consistent with the notion that conoid extrusion serves as a gatekeeper to directs F-actin into the pellicular space, F-actin did not accumulate at the basal pole in cells depleted for MyoA, but instead sequestered at the APR level, where MyoA takes the relay of MyoH (Fig.6b,c). In absence of MyoH, accumulation of F-actin at the basal pole is absent, however due to the block in conoid extrusion, F-actin is seen leaking into the cytosol (Fig.6b,c).

## DISCUSSION

In coccidians, the conoid is topped by PCRs, ring structures that have been well-defined by EM but whose composition and function remain poorly understood ^61^. In this study, U-ExM has unambiguously established that FRM1, GAC and AKMT are positioned at the PCRs, corroborating a previous study that localized AKMT to a ring like structure anterior to the conoid ^62^. These three proteins are instrumental for parasite motility and invasion, with AKMT playing a poorly understood role in the accumulation of GAC at the PCRs ^17^ and in the generation of F-actin flux ^18^. The conoid is known to extrude and retract repeatedly in motile extracellular parasites, a process experimentally induced by calcium ionophores, such as ionomycin ^29^. Combining anti-tubulin antibodies with U-ExM resulted in an accurate and direct visualization of conoid extrusion. When revisiting a series of mutants and pharmacological treatments, it became clear that elongation of extracellular parasites based on phase contrast microscopy ^30^ was not a reliable method to quantify conoid extrusion. While parasites depleted in MyoH were still able to elongate, the conoid remained retracted, a phenotype which was previously overseen ^23^. The 3Me-Pan antibodies that stain both the apical cap and the conoid clearly distinguished conoid extrusion from parasite elongation. It is plausible that lysine methylation modulates actin polymerization by directly modifying either FRM1 or actin. In this context, the staining at the basal pole observed with the 3Me-Pan antibody on motile parasites could correspond to methylated F-actin (Fig.1a). This assay, which cannot be used in case of AKMT depleted parasites, confirmed that conoid movement is dependent on FRM1 and MyoH. The fact that parasites depleted in MyoH, FRM1, AKMT or treated with CD ^23, 24^ were still able to secrete their micronemes indicates that conoid extrusion is not a prerequisite for organelle exocytosis. Conversely, TFP1-depleted parasites in which microneme secretion is abrogated, showed no defect in conoid extrusion or parasite elongation. Of relevance, the signaling cascade leading to conoid extrusion depends on PKG, AKMT and CDPK1, all being required for activation of the actomyosin system. In contrast, remodeling of the parasite cytoskeleton during elongation is independent of conoid extrusion, actin polymerization and microneme secretion but is blocked by C1, an inhibitor of PKG and CDPK1. A change in curvature of the SPMTs has recently been associated with parasite motility ^63^ however the molecular mechanism responsible for this change remains to be discovered. In addition to known invasion factors, we have identified and characterized 5 new components of the PCRs with, Pcr4 and Pcr5 being essential for the integrity of the PCRs. Consistent with a role of PCRs as a hub for actin polymerization, parasites depleted in either Pcr4 or Pcr5 were impaired in conoid extrusion and immotile. Pcr4 and Pcr5 are conserved across Apicomplexa and also form a ring above the conoid in *P. berghei*. The localization of *Pb*GAC and functional dissection of *Pb*Pcr4 and *Pb*Pcr5 indicate conserved roles within the core of PCRs, themselves serving as a platform to assemble components of the glideosome.

Functional and biochemical dissection of Pcr4 and Pcr5 confirmed a similar function and assembly into stable heterodimers *in vitro*. The predicted 3D structure of the Pcr4-Pcr5 heterodimer positions their respective B-Box domains at the extremities of the complex, away from the dimerization interface. Furthermore, mutagenesis of Pcr5 B-Box domains revealed they are dispensable for heterodimerization *in vitro* but instrumental for apical localization of Pcr5 *in vivo*. While Pcr4 and Pcr5 are key components of the PCRs, the structure is predicted to be very large and likely includes additional structural components, themselves dependent on Pcr4 & 2 B-Box domains for PCRs assembly.

The central role played by PCRs towards assembly of glideosome components led to the notion that conoid extrusion serves as gatekeeper for entry of F-actin into the pellicular space to tightly control parasite motility. F-actin is gated into the pellicular space exclusively when the conoid is extruded, where it is translocated to the posterior pole by MyoA. Following invasion and conoid retraction, F-actin no longer enters the pellicular space, which immediately results in halted motility ^64^. The accumulation of F-actin around the conoid in parasites depleted in MyoA but not MyoH supports this model. The mechanism governing extrusion and retraction of the conoid will be of pivotal importance to understand how parasite egress and invasion are controlled.

## Supporting information

Supplementary Discussion

## ACKNOWLEDGMENTS

We thank for their technical assistance, the team at the Bioimaging Core Facility, Rémy Visentin at the Protein platform and Alexandre Hainard at the proteomics platform of the faculty of Medicine University of Geneva as well as Florence Pojer, Luciano Abriata and Kelvin Lau from the “Protein Production and Structure Core Facility” at the EPFL. We thank David Baker for supply of C1. This work was supported by the Swiss National Foundation to D.S.-F. (310030_185325) and to M.B. (BSSGI0_155852 and 310030_208151). L.B. was supported by the Novartis Foundation (19C189). B.M. was supported by the European Research Council (ERC) under the European Union’s Horizon 2020 research and innovation program under grant agreement no. 695596.

## AUTHOR CONTRIBUTIONS

**Conceptualization**, D.S-F., O.V., N.D.S.P., L.B. and M. B. **Methodology**, N.D.S.P., O.V., R.H., L.B., B.M., N.T. **Investigation**, N.D.S.P., O.V., R.H., L.B. and N.T. **Formal Analysis**, N.D.S.P., O.V., R.H., L.B., N.T. and B.M. **Writing – Original Draft**, D.S-F., O.V. and N.D.S.P; **Writing – Review & Editing**, D.S-F., O.V., N.D.S.P, R.H., L.B., N.T., M.B.; **Funding Acquisition** D.S-F. and M.B. **Resources**, D.S-F., O.V. and M.B.; **Supervision** D.S-F., O.V. and M. B. All authors contributed to the article and approved the submitted version.

## COMPETING INTERESTS

The authors declare no competing interests.

## METHODS

### Experimental model and ethic statement

All animal experiments were conducted with the authorization numbers GE-82-15 and GE-41-17, according to the guidelines and regulations issued by the Swiss Federal Veterinary Office.

### Toxoplasma gondii maintenance and transfection

*Toxoplasma gondii* tachyzoites were amplified in human foreskin fibroblasts (HFFs, ATCC) in Dulbecco’s Modified Eagle’s Medium (DMEM, Gibco) supplemented with 5% of Fetal Bovine Serum (FCS, Gibco), 2 mM glutamine and 25 µg/ml gentamicin (Gibco).

Tachyzoites were transfected by electroporation ^65^. To target any gene of interest, 40 µg of specific gRNA was transfected alongside a PCR product flanked by homology regions. The list of primers to generate the gRNAs and the PCR products are listed in the Supplementary Table 2. Parasites carrying an HXGPRT cassette were selected with 25 mg/ml of mycophenolic acid and 50 mg/ml of xanthine. Parasites carrying a DHFR cassette were selected with 1 µg/ml of pyrimethamine. In the case of second copy genes inserted in the UPRT locus, the parasites were selected with 5 µM of 5-fluorodeoxyuridine (FUDR).

Auxin-degron fused proteins depletions (AiD strains) were achieved by addition of 500 µM of auxin (IAA) ^66^. Depletion via the Tet-inducible system (iKD strains) was achieved by addition of 1 µg/ml of anhydrotetracycline (ATc) ^67^. The list of strains generated/used in this study is presented in the Supplementary Table 2.

### Plasmodium berghei maintenance and transfection

Parental cell lines were derived from *P. berghei* ANKA strain-derived clone 2.34 ^68^ in which the Celtos ^69^ is tagged at the endogenous locus with triple HA tag. Parental cell lines together with derived transgenic lines were grown and maintained in CD1 outbred mice. Six to ten weeks-old mice were obtained from Charles River laboratories and females were used for all experiments. Mice were specific pathogen free (including *Mycoplasma pulmonis*) and subjected to regular pathogen monitoring by sentinel screening.

They were housed in individually ventilated cages furnished with a cardboard mouse house and Nestlet, maintained at 21 ± 2°C under a 12 hr light/dark cycle and given commercially prepared autoclaved dry rodent diet and water ad libitum. The parasitaemia of infected animals was determined by microscopy of methanol-fixed and Giemsa-stained thin blood smears. For gametocyte production, parasites were grown in mice that had been phenyl hydrazine treated three days before infection. Exflagellation was induced in exflagellation medium (RPMI 1640 containing 25 mM HEPES, 4 mM sodium bicarbonate, 5% fetal calf serum (FCS), 100 mM xanthurenic acid, pH 7.8). For gametocyte purification, parasites were harvested in suspended animation medium (SA; RPMI 1640 containing 25 mM HEPES, 5% FCS, 4 mM sodium bicarbonate, pH 7.20) and separated from uninfected erythrocytes on a Histodenz cushion made from 48% of a Histodenz stock (27.6% [w/v] Histodenz [Sigma/ Alere Technologies] in 5.0 mM TrisHCl, 3.0 mM KCl, 0.3 mM EDTA, pH 7.20) and 52% SA, final pH 7.2. Gametocytes were harvested from the interface.

Schizonts for transfection were purified from overnight *in vitro* culture on a Histodenz cushion made from 55% of the Histodenz stock and 45% PBS. Parasites were harvested from the interface and collected by centrifugation at 500 g for 3 min, resuspended in 25 µl Amaxa Basic Parasite Nucleofector solution (Lonza) and added to 10 µg DNA dissolved in 10 µl H2O. Cells were electroporated using the FI-115 program of the Amaxa Nucleofector 4D. Transfected parasites were resuspended in 200 ml fresh RBCs and injected intraperitoneally into mice. Parasite selection with 0.07 mg/mL pyrimethamine (Sigma) in the drinking water (pH ∼4.5) was initiated one day after infection.

### Generation of transgenic parasites targeting constructs for Plasmodium

The oligonucleotides used to generate transgenic parasite lines are in Supplementary Table 2. C-terminal tagging of *P. berghei* proteins by pCP was as described ^70^ and targeted endogenous loci by allele replacement. Briefly sequences comprising ∼ 500 bp from the C-terminus of the coding sequence and ∼500 bp from the immediate 3’ UTR for genes Pcr4 and Pcr5 were cloned into KpnI and XhoI sites upstream to the coding sequences of mNeonGreen fused to triple hemagglutinin epitope tag in pCP-mNG-3xHA, along with a NotI linearization site between the targeting sequences. For stage-specific knockdown of Pcr4 and Pcr5, constructs were iterated from pCP, described previously, for easier “one-step” plasmid generation and to integrate a 3xHA tag at the N-terminus of the target endogenous locus by allele replacement. Briefly, the promoter sequence for *clag9* ^71^ and coding sequence for 3xHA were amplified using primers MB955 with MB956 and MB1048 with MB1049, respectively, and replaced the corresponding fragment between ApaI and KpnI sites in pCP, separated by a XhoI site between the sequences. Sequences comprising ∼ 500 bp from the N-terminus of the coding sequence and ∼500 bp from the immediate 5’ UTR for genes Pcr4 and Pcr5 were cloned into XhoI and EcoRI sites downstream to the coding sequences of 3xHA, along with a NotI linearisation site between the targeting sequences.

### Ultrastructure-Expansion Microscopy (U-ExM)

For U-ExM, the same general protocol was used for both *Plasmodium* and *Toxoplasma* ^36^. In short, parasites were resuspended in warm PBS (with BIPPO in the case of *Toxoplasma*) and sedimented on coverslips coated with poly-lysine (Gibco). The excess liquid was then discarded and the coverslips were incubated for 5h at 37°C in a mix of 0.7% formaldehyde / 1% acrylamide. The coverslips were then used to cover drops of monomer solution (19% sodium acrylate / 10% acrylamide / 0.1% bis-acrylamide) supplemented with 0.5% ammonium persulfate and 0.5% tetramethylethylenediamine (TEMED) to initiate the gel polymerization. After an incubation of 30 minutes at 37°C, the fully polymerized gels were detached from the coverslip and incubated 1h30 at 95°C in denaturation buffer (200mM SDS, 200mM NaCl, 50 mM Tris in water, pH 9). Gels were then expanded overnight in water.

The following day, the days the expansion ratio of each gel was determined by measuring their diameter (here all the gels presented had an expansion ratio of 4). Then, the gels were shrinked in PBS, cut, and incubated for 3h at 37°C with agitation in a mix of appropriate primary antibody diluted in freshly prepared PBS-BSA 2%. The list of antibodies used and their dilution is listed in the Key Resource Table. After 3 washes of 10 minutes in PBS-Tween 0.1%, the gels were incubated 2h at 37°C (with agitation and in the dark) in a mix of appropriate secondary antibodies. After 3 washes of 10 minutes in PBS-Tween 0.1%, the gels were expanded overnight in water. Imaging was performed using Leica TCS SP8 inverted microscope, equipped with an environmental chamber and an “IceCUBE” temperature control system (Life Imaging Services). The microscope was used with an HC PL Apo 100x/ 1.40 Oil CS2 objective and with HyD detectors. Pieces of gels were cut and placed on a 24mm poly-lysine coated coverslip (to avoid gel drifting during acquisition) fitted in a metallic O-ring 35mm imaging chamber (Okolab). Z-stack were acquired with the Leica LAS X software and deconvolved with the built-in “Lightning” mode. Images were then processed with the ImageJ software and maximum projections were used for publication.

### Conoid extrusion assay by U-ExM

For *Toxoplasma*, conoid extrusion was induced using 10 µM of BIPPO. For each biological replicate (n=3), two pieces of the same gel were used, and approximately 100 parasites were counted in each piece. C1 corresponds to 4-[2-(4-Fluorphenyl)-5-(1-methylpiperidine-4-yl)-1*H* pyrrol-3-yl] pyridine) ^72^. For *Plasmodium* ookinetes, only gels with an expansion ratio of 4 were used. Between 25 and 30 ookinetes were assessed in each biological replicate. A single picture (1 single stack) was acquired for each ookinete. Only ookinetes lying perfectly on their side were used. The resulting midsagittal picture of the apical complex was then used to measure the distance between the APR and the conoid with the Leica LAS X software.

### Extrusion/protrusion assay by IFA for Toxoplasma

Freshly egressed or syringe-lysed tachyzoites were incubated with the appropriate drug (2µM cytochalasin D for 30min; 1µM compound 1 for 30min). Parasites were then seeded on gelatin-coated coverslips, and warm media containing BIPPO was added on top. Coverslips were centrifuged at 1000rpm for 1 minute, and incubated 10 minutes at 37°C. Parasites were then fixed and stained, with anti-actin and anti-methylated lysine antibodies. For each condition, 200 parasites were counted, for each of the three independent biological replicates (n=3).

### Immunofluorescence assay for Toxoplasma

HFF seeded on coverslips were infected with T. gondii tachyzoites and let grow at 37°C for 15h-24h. Parasites were then fixed with 4% PFA/0.05% glutaraldehyde (PFA-GA) in PBS or ice-cold methanol, neutralized in 0.1M glycine-PBS for 3–5 min in the case of PAF-GA fixation and processed as previously described ^73^. Images were obtained with a Zeiss laser scanning confocal microscope (LSM700 using objective apochromat 63x/1.4 oil). Images were then processed for publication with the ImageJ software.

### Protein localization for Plasmodium

For localization of tagged proteins in *P. berghei* by native fluorescence, cells from different proliferative stages during the parasite lifecycle were mounted in exflagellation medium (RPMI 1640 containing 25 mM HEPES, 4 mM sodium bicarbonate, 5% fetal calf serum (FCS), 100 mM xanthurenic acid, pH 7.8), and imaged using an inverted Zeiss Axio Observer Z1 microscope fitted with an Axiocam 506 mono 14 bit camera and Plan Apochromat 63x / 1.4 Oil DIC III objective. All images of fluorescent proteins were captured at RT with equal exposure settings. Images for level comparison were processed with the same alterations to minimum and maximum display levels. Analysis and displays were performed in Fiji and the statistical programming package R (http://www.r-project.org).

### Plaque assay

HFF monolayer were infected with freshly egressed parasites and grown for 7 days at 37°C. Cells were then fixed with PFA-GA, subsequently neutralized in 0.1M glycine/PBS. The cells were then washed in PBS and stained with crystal violet.

### Invasion assay

Coverslips covered with HFF monolayer were infected with T. gondii tachyzoites, centrifuged at 1000rpm for 1 minute and incubated at 37°C for 30 minutes. Cells were then fixed with PFA-GA. Fixed cells were then incubated in 5% PBS-BSA for 20 minutes of blocking, 1h with anti-SAG1 antibody and washed 3 times in PBS. Cells and antibodies were then fixed with 1% formaldehyde for 7 min, permeabilized with 0.2% Triton X100 in PBS for 20 minutes. Finally, cells were incubated with anti-GAP45 antibodies, washed 3 times, and incubated with appropriate secondary antibodies. 200 parasites were counted for each condition and over three independent biological replicates (n=3).

### Induced egress assay

Coverslips covered with HFF monolayer were infected with T. gondii tachyzoites, centrifuged at 1000rpm for 1 minute and incubated at 37°C for 30 hours. Cells were then incubated with DMEM media containing BIPPO for 10 minutes at 37°C. Cells were then fixed with PFA-GA labelled as described previously, with anti-GAP45 and anti-GRA3 antibodies. 200 vacuoles were counted for each condition and over three independent biological replicates (n=3).

### Gliding (trail deposition) assay for Toxoplasma

Freshly egressed tachyzoites were resuspended in warm DMEM media containing BIPPO. Parasites were then put on poly-lysine coated coverslips, centrifuged at 1000rpm for 1 minute, and incubated 10 minutes at 37°C. Parasites were then fixed and stained, as described previously, with anti-SAG1 antibodies (and in absence on Triton X100).

### Microneme secretion

Freshly egressed parasites (syringed out in the case of parasite with severe egress defects) were washed twice in warm DMEM media. Parasites pellets were then resuspended in media containing BIPPO and incubated at 37°C for 10 minutes. Pellets and supernatant (ESA) were then separated by centrifugation at 2000 x g. The ESA fraction was then centrifuged at 5000 x g to remove any cells debris. Pellets fractions were washed once in PBS to remove any ESA remaining. Pellets and ESA were then analyzed by western blot using anti-MIC2, anti-Catalase, and anti-GRA1 antibodies. Experiments were performed three times independently (n=3) and a representative replicate is presented in the manuscript.

### Second copy expression of Pcr5 at the UPRT locus

A codon-remodeled copy of the Pcr5, with the B-Box domains flanked by MfeI and PstI sites, inserted in a pUC57-Kan vector (PvuII sites) was ordered from GENEWIZ®. Pcr5 was excised from the pUC57-Kan vector (using PvuII) and inserted in a linearized (SfoI and SmaI) pUC19-Amp vector. Good integration was checked by PCR.

In parallel, gBlocks® Gene Fragment containing the three versions of the mutated Bbox (MUT1, MUT2 and MUT3), flanked by EcoRI and NsiI sites, were ordered from IDT®. All three were independently inserted in a pCR-BluntII vector via TOPO reaction. Good integration was checked by PCR. Final constructs were checked by sequencing.

pUC19-Pcr5-Amp was digested with MfeI and PstI while all three Bbox mutant construct were digested with EcoRI and NsiI enzymes and purified before ligation. The three resulting pUC19-Pcr5MUTx-Amp vectors were then checked by sequencing.

The three versions of Pcr5 (Pcr5MUT1, Pcr5MUT2 and Pcr5MUT3) were finally excised from the pUC19 vector (MfeI/PstI) and inserted in the pUPRT-pTUB-G13-Ty vector (the G13 gene being excised with EcoRI/PstI) ^74^. Good integration was checked by PCR. Final constructs were checked by sequencing.

For transfection, 60µg of pUPRT-pTUB-Pcr5MUTx-Ty vector were linearized by NdeI and mixed with 20µg of gRNA targeting the UPRT locus.

### Protein cloning for expression in insect cells

Recombinant Pcr5 alone or in complex with Pcr4 were purified following a similar protocol. Pcr4 and Pcr5 coding sequences were each inserted into pFastBac vectors encoding a C-terminal 10-His tag and a C-terminal twin-Strep tag respectively. Baculoviruses encoding the desired proteins were generated following bac-to-bac standard procedures (Invitrogen). Protein expression was performed in baculovirus-infected Sf9 cells for 68 h at 27°C before harvesting and purification.

### Pull down and Solubility Assays insect-cell expressed PCR proteins

Sf9 cells were co-infected with baculoviruses (1/100e dilution for each expressing Pcr4 and Pcr5 (WT, Mut1, Mut2, Mut3) during 68 h at 27°C in a final volume of 15 mL. Cells were harvested and the pellet were resuspended in 6 mL of Lysis Buffer (50 mM Tris-HCL pH 7.5, 800 mM NaCl, 4 mM 2-Mercaptoethanol). Cells were then lysed by sonication (10 shoots, 60% output control, 20% duty cycle) and cell lysates were centrifuged at 35,000 × g for 30min at 4°C. To check the expression and the solubility of each single and co-infection 25µL of the cell lysate and supernatant was diluted in 5µL of SDS-blue 6X. To assess the interaction between Pcr4 and Pcr5, the supernatants were applied to 1mL Strep-trap column (GE healthcare) and His-trap column (GE healthcare) previously equilibrated with Lysis Buffer. Columns were washed with 5mL of Lysis Buffer and the protein was eluted in 3mL of Elution Buffer. Elution Buffer for Strep-trap Columns was made with Lysis Buffer supplemented with 1mg/mL of Biotin. Elution Buffer for His-trap Columns was made with Lysis Buffer supplemented with 400mM of Imidazole. Eluted complexes were concentrated around 50µL (Vivaspin 200µL) and diluted in 10µL of SDS-blue 6X. Samples were loaded on SDS-PAGE colored with Coomassie Blue (Lubio Science).

Protein solubility for both PCR1 and PCR2 was assessed by band densitometry on coomassie-stained gels. Lysates and supernatants of Sf9 insect cells overexpressing the PCR proteins of interest were compared to lysates and supernatants obtained from non-infected cells. To support the band densitometry data, the same samples were quantified by western blot using antibodies specifically detecting the tags attached to either PCR1 (His-tag) or PCR2 (STREP-tag).

### Hydrogen/ Deuterium exchange coupled to Mass Spectrometry (HDX-MS)

HDX-MS experiments were performed at the UniGe Protein Platform (University of Gene-va, Switzerland) following a well-established protocol with minimal modifications ^75^. HDX reactions were done in 50 μl volumes using 380 pmol (7.6 μM) of recombinant purified Pcr4-2 complex. The same procedure was applied to both Pcr4-2 complexes: WT and Mut1. Briefly, 12 μl of protein complex was preincubated at RT before addition of 38 μl of deuterated buffer (20 mM Tris pH 7.5, 800 mM NaCl in D_2_O). Reactions were carried-out for 3, 30 and 300 sec at room temperature, and terminated by the sequential addition of 20 μl of icecold quenching buffer (4M Guanidine-HCl/ 1M NaCl/ 0.1M NaH_2_PO_4_ pH 2.5/ 1% formic acid). Samples were immediately frozen in liquid nitrogen and stored at -80 °C for up to one week. To quantify deuterium uptake, protein samples were thawed and injected in UPLC system immersed in ice. The protein was digested via two immobilized pepsin columns (Thermo #23131), and peptides were collected onto a VanGuard pre-column trap (Waters). The trap was subsequently eluted and peptides separated with a C18, 300 Å, 1.7 μm particle size Fortis Bio column 100 × 2.1 mm over a gradient of 8 – 30 % buffer B over 20 min at 150 μl/ min (Buffer A: 0.1% formic acid; buffer B: 100% acetonitrile). Mass spectra were acquired on an Orbitrap Velos Pro (Thermoscientific), for ions from 400 to 2200 m/z using an electrospray ionization source operated at 270°C, 5 kV of ion spray voltage. Peptides identified by data-dependent acquisition after MS/MS and data were analyzed by Mascot. A 73 % and 69 % sequence coverage for Pcr5 and Pcr4 respectively was obtained. Deuterium incorporation levels were quantified using HD examiner software (Sierra Analytics), and quality of every peptide was checked manually. Results are presented as percentage of theoretical maximal deuteration level. All experimental details and data of percentage deuterium incorporation for all peptides can be found in Supplementary Tables 1.

### Electron microscopy

As described previously ^52^, extracellular parasites were pelleted in PBS. Conoid protrusion was induced by incubation with 40 µL of BIPPO in PBS for 5 min at 37°C. 4 µL of the sample were applied on glow-discharged 200-mesh Cu electron microscopy grid for 10 min. The excess of the sample was removed by blotting with filter paper and immediately washed 3 times on drop of double distilled water. Finally, the sample was negatively stained with 0.5% aqueous solution of phosphotungstic acid (PTA) for 20 s and air dried. Electron micrographs of parasites apical poles were collected with Tecnai 20 TEM (FEI, Netherland) operated at 80 kV acceleration voltage equipped with side mounted CCD camera (MegaView III, Olympus Imaging Systems) controlled by iTEM software (Olympus Imaging Systems).

### Exflagellation analysis

The exflagellation rate was assessed by observing exflagellation centers following induction under a light microscope with a 40x objective (n=3).

### Ookinete conversion

For ookinetes, gametocyte-rich mouse blood was inoculated with 9 volumes of ookinete medium and incubated for 18-20 hr at 19°C. The conversion efficiency was determined by adding anti-p28-Cy3 monoclonal antibody and calculating the ratio of mature banana-shaped p28-positive ookinetes compared to round cells and developmentally arrested zygotes (n=3).

### Motility assay for Plasmodium ookinetes

Ookinete motility was assessed as described previously ^76^ in Matrigel (BD Biosciences). Time-lapse videos of ookinetes were acquired on an inverted Zeiss Axio Observer Z1 microscope fitted with an Axiocam 506 mono 14 bit camera and Plan Apochromat 63x / 1.4 Oil DIC III objective. Tracking of the ookinetes was performed with the “Chemotaxis and Migration Tool” available online (https://ibidi.com/chemotaxis-analysis/171-chemotaxis-and-migration-tool.html). The gliding speed was determined by quantifying the total covered distance. Videos were acquired at 3 frames/min over the course of 10 minutes (n=3).

### Bioinformatic analyses

Searches were as described ^70^ and based on predicted protein datasets for 90 alveolate organisms from genomes and transcriptomes. For transcriptomes, ORFs were predicted using TransDecoder. Apicomplexan orthologous groups were generated and defined by OrthoFinder ^77^. Orthogroups including PCR proteins were aligned using MAFFT ^78^, the L-INS-i strategy, trimmed to conserved regions with trimAl, modelled by HMMER3 ^79^ and searched to find similar sequences in all predicted proteomes. Hits were added iteratively and used to create new profiles until no new sequences were identified. Alignments were visualized and modified using Jalview ^80^. Profile–profile comparisons were performed using HH-suite3 ^81^. For phylogenies, trimmed alignments were used to infer maximum likelihood phylogenies using IQ-TREE2 ^82^. Trees were visualized in FigTree (tree.bio.ed.ac.uk/software/figtree) and Phytools ^83^, as part of the statistical programming package ‘R’.

## Data availability

All data are available within the paper and its supplementary information. Sequences used in this study have been obtained from EuPathDB. HDX-MS data are available via ProteomeXchange with identifier PXD031816. All biological materials and data are available from the author upon request.

---

**Extended Figure 1.**
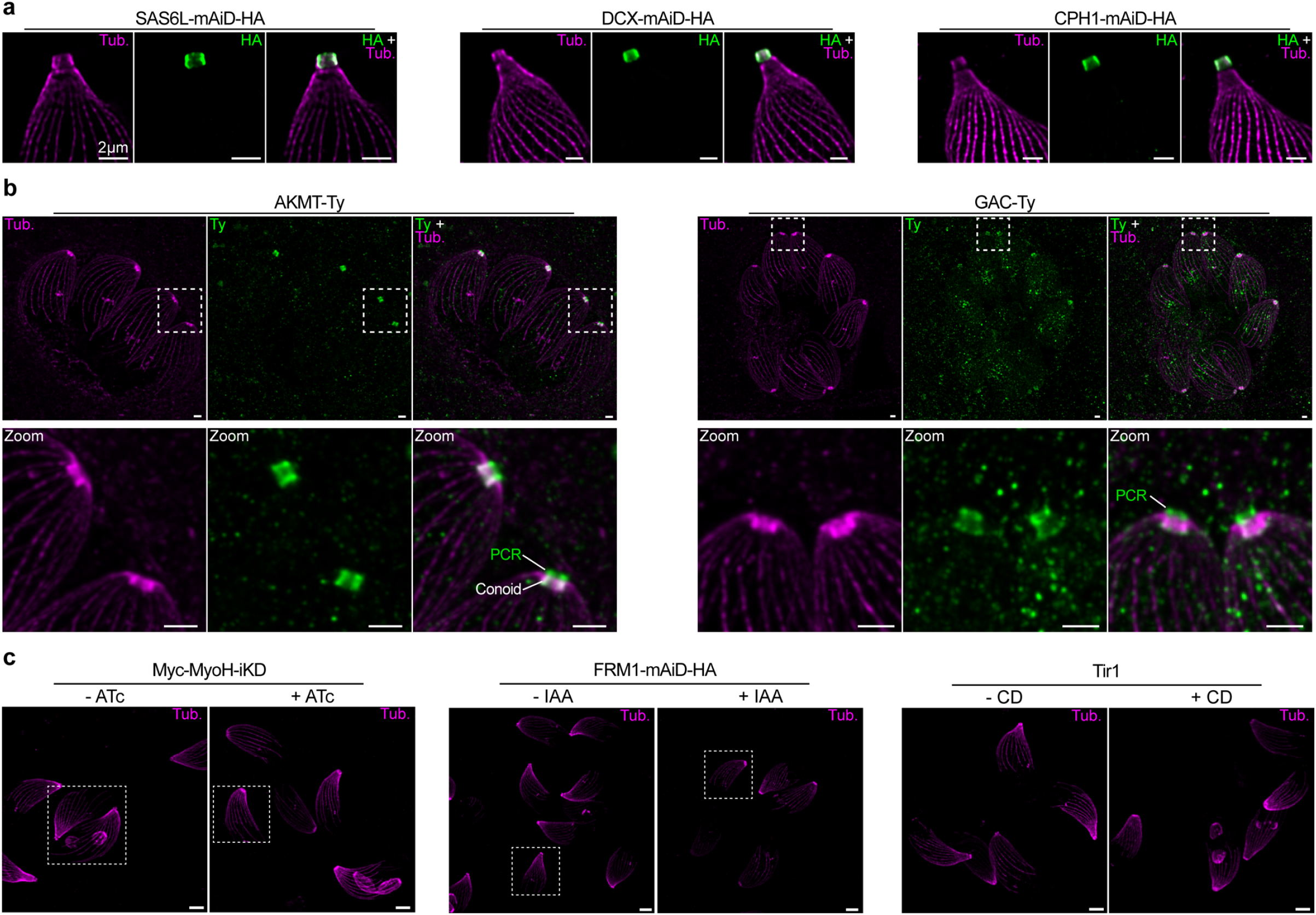
**(a)** U-ExM localization of conoidal proteins fused with a mAiD-HA cassette. Scale bar = 2µm. **(b)** Localization of AKMT and GAC in intracellular vacuoles as seen by U-ExM. Scale bar = 2µm. **(c)** Representative pictures of some U-ExM samples used for quantification of Figure 1C. Dash boxes represents crop parasites displayed in Fig.1b. Scale bar = 5µm.

**Extended Figure 2.**
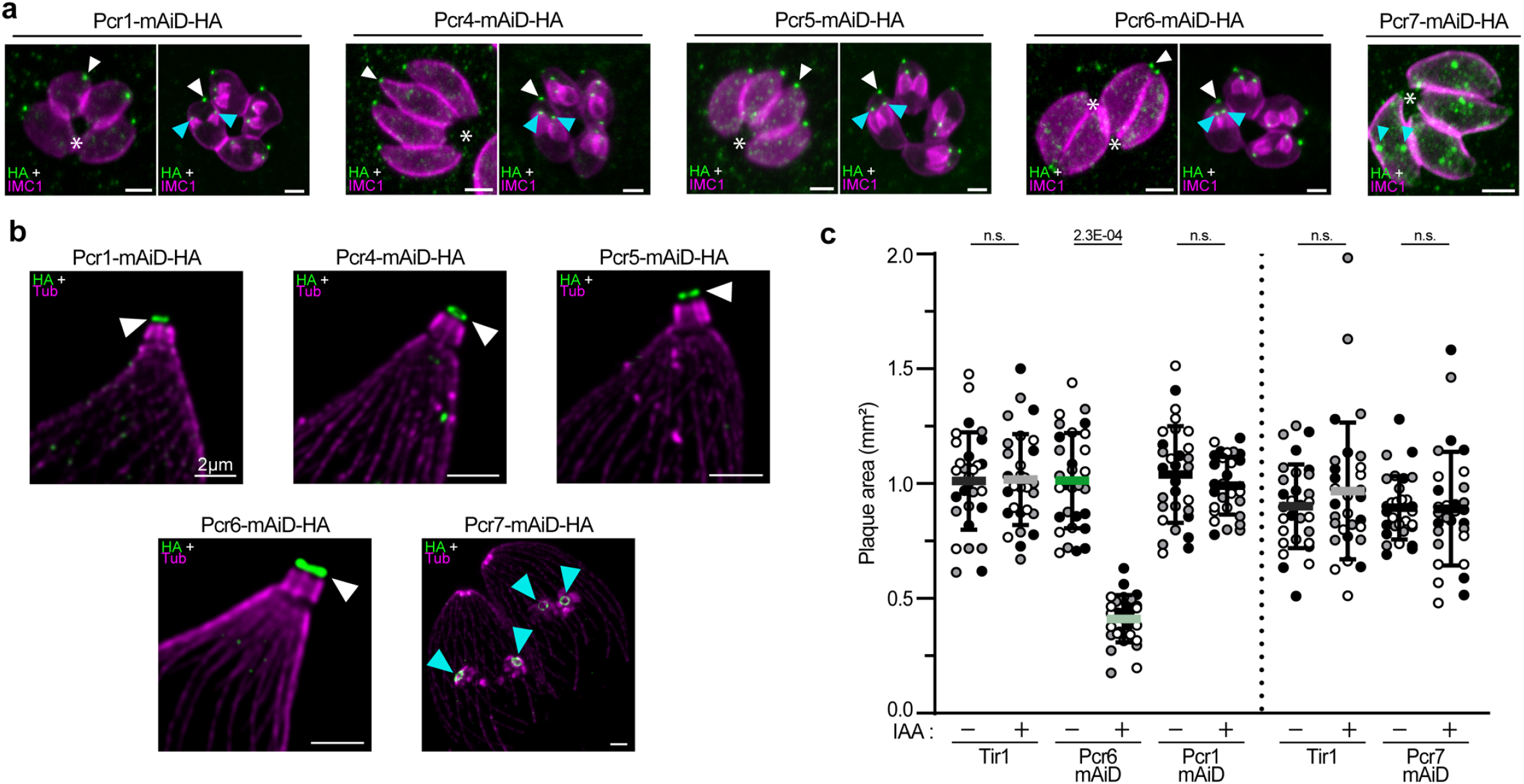
**(a)** IFA of Pcr1-4-5-6-7 in non-dividing and dividing parasite showing that Pcr1-4-5-6 are also found in forming daughter cells. White arrowhead = apical pole of mature cells. Cyan arrowhead = apical pole of forming daughter cells. Scale bar = 2µm. **(b)** Additional U-ExM pictures of Pcr1-4-5-6-7. Scale bar = 2µm. **(c)** Quantification of the plaque size. The different replicates are found in shades of grey. Ten plaques were measures for each condition and replicate (n=3). Unpaired t-test where non-significant (n.s.) if P>5E-02.

**Extended Figure 3.**
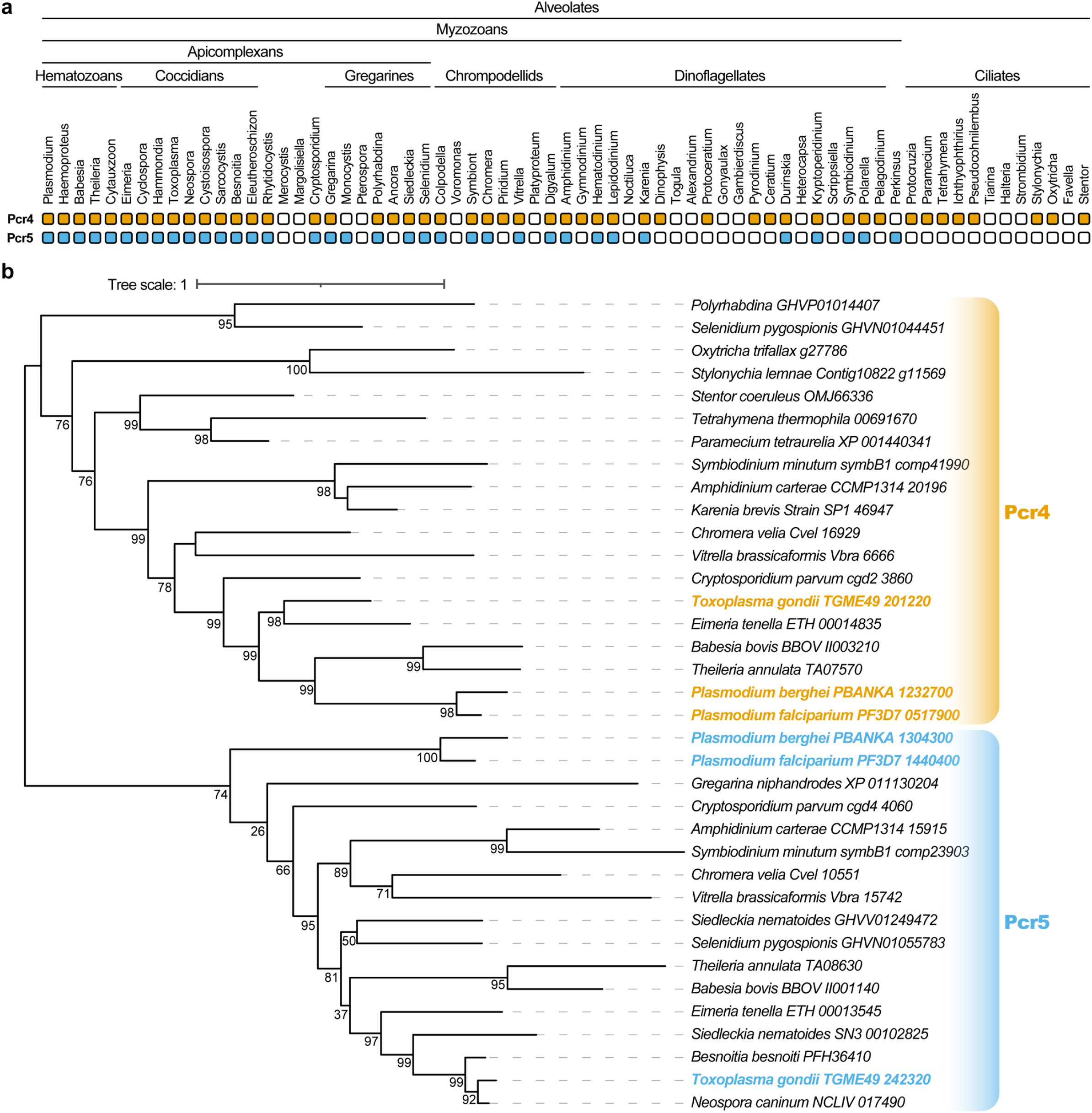
**(a)** Conservation of Pcr4 and Pcr5 in the Alveolate superphylum. Presence (coloured) or absence (white) of homologs identified across alveolate predicted proteomes following. **(b)** Result of a maximum likelihood inference based on an alignment of sequences retrieved for homologs of Pcr4 and Pcr5. Numbers beside nodes indicate bootstrap support (1000 ultrafast bootstrap replicates).

**Extended Figure 4.**
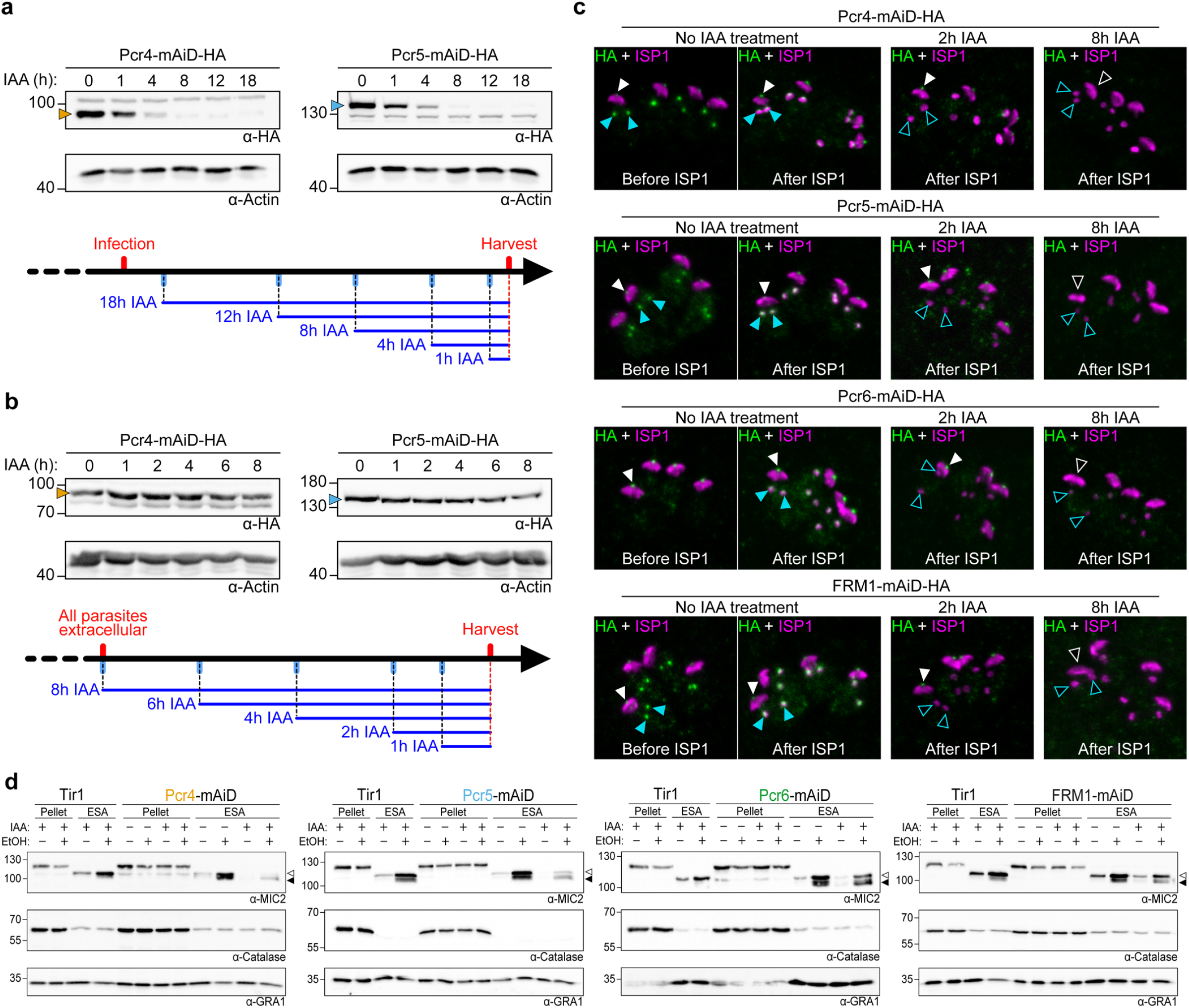
**(a)** Regulation of Pcr4 and Pcr5 on intracellular parasites. Time of auxin incubations are indicated in hour. Actin is used as loading control. A small scheme of the experimental timeline is presented. **(b)** Regulation of Pcr4 and Pcr5 on extracellular parasites. Time of auxin incubations are indicated in hour. Actin is used as loading control. A small scheme of the experimental timeline is presented. **(c)** Stability of Pcr4-5-6 and FRM1 fused with a mAiD-HA cassette, after short IAA treatment. ISP1 is one the earliest marker appearing during daughter cells biogenesis. White arrowhead = apical signal at the apical pole of mature cells. Cyan arrowhead = apical signal at the apical pole of forming daughter cells. Black filled arrowhead = absence of signal at the apical pole. **(d)** Representative pictures of microneme secretion quantified in Fig.1d. White arrowhead = secreted MIC2. Black arrowhead = secreted MIC2 trimmed by SUB1 protease. GRA1 is used as loading control while catalase is used as a lysis control.

**Extended Figure 5.**
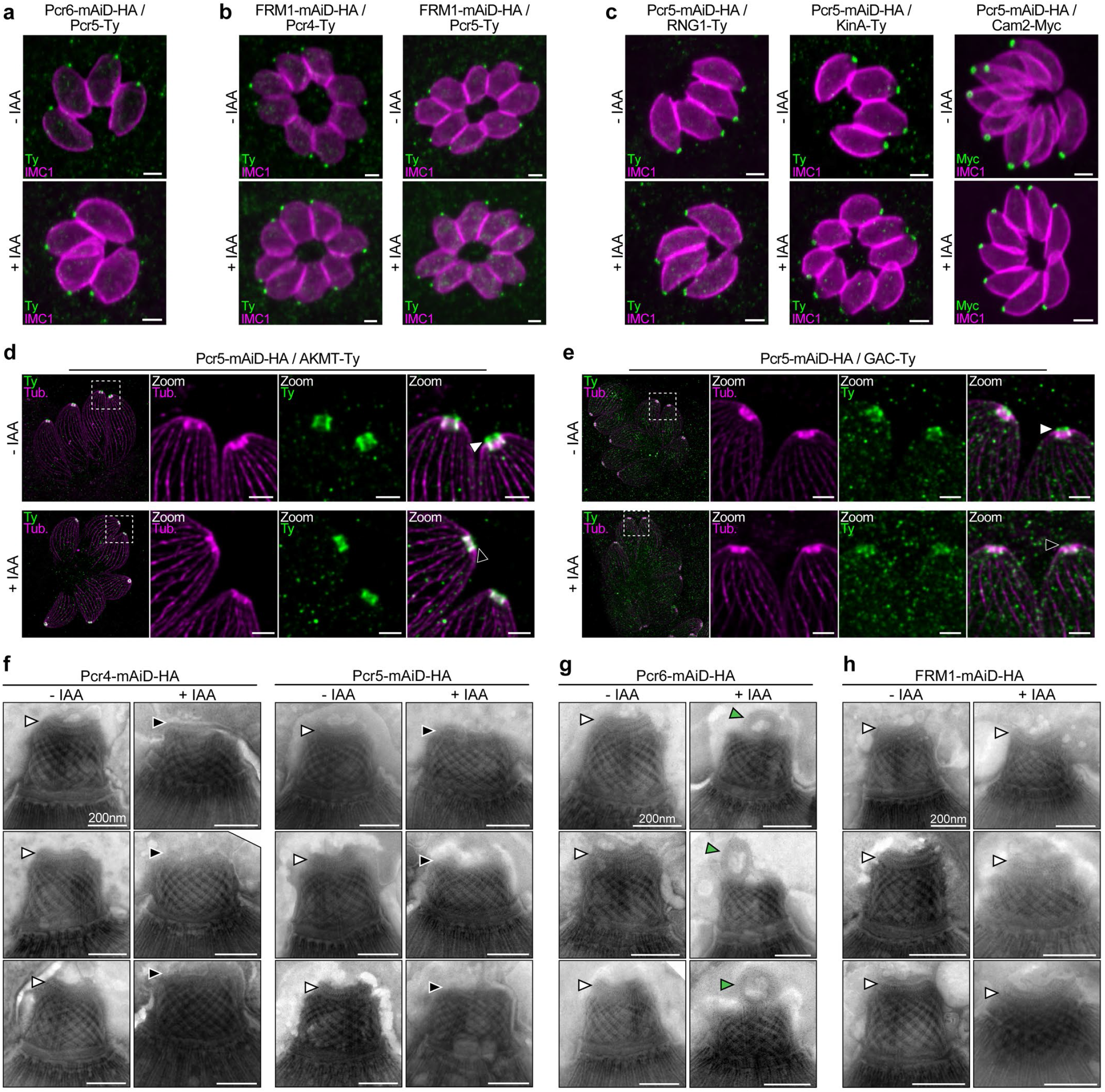
**(a)** IFA showing that the depletion of Pcr6 did not induce the loss of Pcr5. Scale bars = 2 µm. **(b)** IFA showing that the depletion of FRM1 did not induce the loss of Pcr4 and Pcr5. Scale bars = 2 µm. **(c)** IFA showing that the depletion of Pcr5 does not affect conoid (Cam2) and APR (RNG1/KinA) markers. **(d)** Depletion of Pcr5 leads to the loss of the PCRs signal of AKMT in intracellular parasite assessed by U-ExM. Scale bars = 2 µm. **(e)** Depletion of Pcr5 leads to the loss of the PCRs signal of GAC in intracellular parasite assessed by U-ExM. Scale bars = 2 µm. **(f)** Gallery of additional EM images of Pcr4 and Pcr5 knock-down strains. Scale bar = 200 nm. **(g)** Gallery of additional EM images of Pcr6 knock-down strains. Scale bar = 200 nm. **(h)** Gallery of EM images of FRM1 knock-down strains. Scale bar = 200 nm.

**Extended Figure 6.**
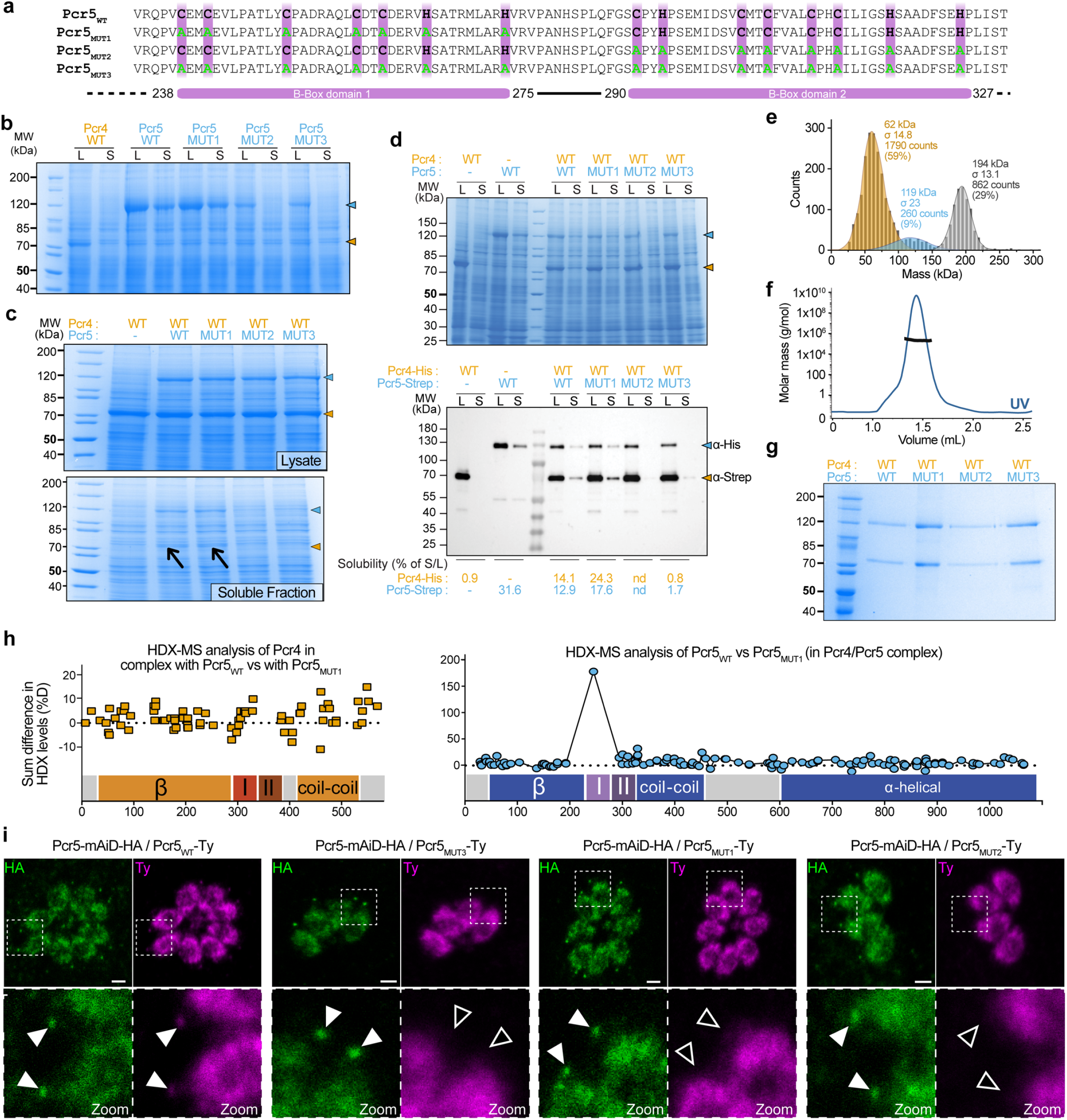
**(a)** Schematic of Pcr5 two B-Box domains. The amino acid sequences of the four versions of Pcr5 used for complementation are presented. Cysteine and histidine residues (violet), known to chelate zinc ions, were mutated to alanine to create mutated versions of the Bbox domains. **(b)** Representative Coomassie stained gel of the solubility assay for the individual proteins quantified in Fig.4a. L = Whole cell lysate. S = Soluble fraction. **(c)** Representative Coomassie stained gel of a solubility assay when Pcr4 is co-expressed with the four versions of Pcr5, quantified in Fig.4b. **(d)** Representative Coomassie stained gels used for solubility quantification and corresponding quantified Western Blot by band densitometry. **(e)** Mass photometry of the Pcr4-Pcr5 complex allowed the measure of the complex mass. **(f)** SEC-MALS coupled with size-exclusion chromatography for the Pcr4-Pcr5 complex. **(g)** Representative Coomassie stained gel of pull-down assays performed with Pcr4 coexpressed with the four versions of Pcr5. Pull-down was performed using anti-Strep column (Pcr5 pull-down). **(h)** HDX-MS analysis highlight a different dynamic of Pcr5 when mutated in the first B-Box domain (Pcr5_MUT1_) **(i)** Immunofluorescence of the four complemented strains. The endogenous copy is presented in green (HA) and the second copy is presented in magenta (Ty). White arrowhead = presence of apical signal. Black arrowhead = no apical signal. Scale bars = 2 µm.

**Extended Figure 7.**
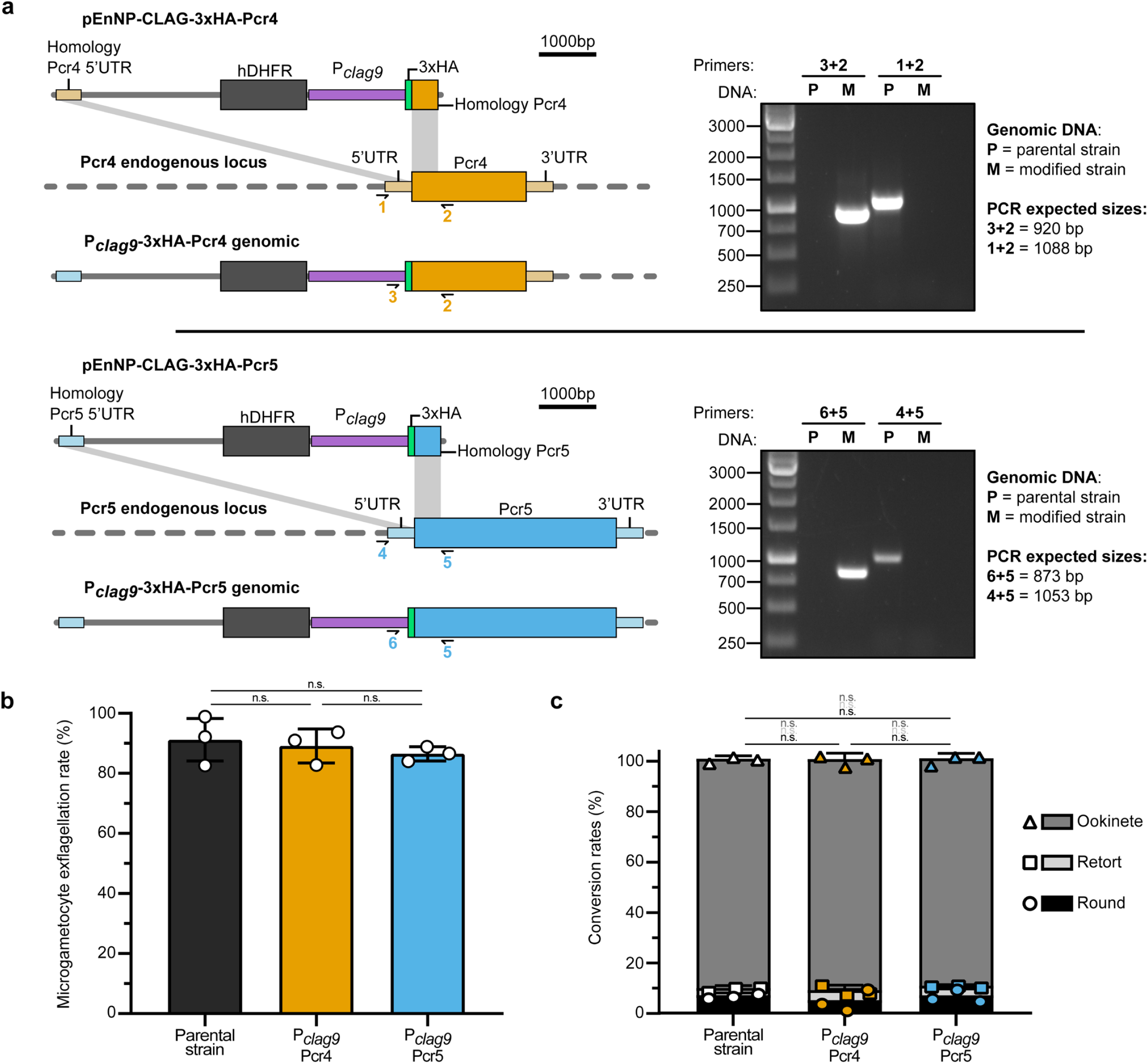
**(a)** Schematics of the promoter-swap strategy to obtain the P_*clag9*_Pcr4 and P_*clag9*_Pcr5 strains. On the right, integration PCR are presented. **(b)** Quantification of microgametocyte exflagellation for the P_*clag9*_Pcr4 and P_*clag9*_Pcr5 strains. Data are presented as mean ± SD (individual biological replicates are also presented). **(c)** Quantification of ookinete conversion for the P_*clag9*_Pcr4 and P_*clag9*_Pcr5 strains. All three states of the conversion (round, retort and ookinete) are presented. Data are presented as mean ± SD (individual biological replicates are also presented). For all statistical analysis, an unpaired t-test where non-significant (n.s.) if P>5E-02 was used. All data are presented as mean ± SD (n=3).

## Notes

### Competing Interest Statement

The authors have declared no competing interest.

