## Supplementary Discussion for "Conoid extrusion serves as gatekeeper for entry of glideosome components into the pellicular space to control motility and invasion in Apicomplexa"

*Repertoire of conoid proteins*

Sub-cellular fractionation studies referred to as Localization of Organelle Proteins by Isotope Tagging (*LOPIT*), have highlighted numerous apical complex proteins, some of which were discernibly validated by high resolution confocal microscopy to either the cone of the conoid, its base, the PCRs or the APR [1]. Most accurate descriptions of the positioning of proteins to specific sub-compartments of the apical complex have been obtained by immuno-EM, revealing the localization of CEN2 at PCRs [2], ICMAP1 at ICMTs [3], and DCX at the cone of conoid [4]. Recently, near-native expansion of biological samples by U-ExM has offered a greatly simplified procedure to localize novel proteins to substructures of the conoid in apicomplexan parasites [5-7].

*Conoid extrusion and F-actin flux*

Depletion of *T. gondii* FRM1 and MyoH, as well as the use of CD, abolished conoid extrusion, making F-actin a central player in this process (Fig.6c). Absence of Pcr4 or Pcr5 led to the disappearance of the PCRs and concomitantly the loss of FRM1, therefore resulting in a defect in conoid extrusion. Importantly, FRM1, not only promotes conoid extrusion powered by MyoH, but also produces F-actin at the origin of the apico-basal flux generated by MyoA.

In contrast, Pcr6 depletion led to a detachment of the PCRs from the conoid, leading to an intermediate phenotype and only partial defect in conoid extrusion. Intriguingly, these parasites only display circular trails when gliding on a 2D surface (Fig.2e). This data support the idea that the F-actin flux in the Pcr6 lacking parasites is abnormal. Indeed, with an abnormal F-actin flux, the secreted adhesins are not translocated efficiently toward the basal pole of the parasite, generating a gradient of adhesins toward the apical pole. While the twirling and helical motility require the apical pole of the parasite to detach from the substrate at some point, circular motility is performed with the apical pole in constant contact with the substrate. It is therefore expected for parasites with a strong adhesion at the apical pole to mostly engage in circular motility. Relevantly, the circular gliding phenotype has been described in other mutants for which adhesins are not efficiently translocated to the basal pole and accumulates at the apical pole, like in the case of MyoA- or MLC1-depleted parasites [8].

The irreversible inhibitor 6 (also referred to as conoidin A), was the only compound identified in a pioneering screen of small molecules interfering with *T. gondii* invasion, to block parasite elongation [9]. Intriguingly, inhibitor 6 had no effect on either parasite motility or microneme secretion and hence is unlikely to block conoid extrusion.

*Preconoidal rings in malaria parasites*

Placing either *Pb*Pcr4 or *Pb*Pcr5 under blood-stage specific expression reduced parasite motility and the distance between the conoid and APR in ookinetes, providing evidence of conoid dynamics dependent on the PCRs in malaria parasites. However, it is more likely that this dynamic is more reminiscent to the "elongation" observed in *Toxoplasma* rather than the "extrusion" of the conoid through the APR. *Pb*MyoB is the closest homolog to *Tg*MyoH, and was also found at the conoid of *P. berghei*, although its role in conoid movement remains to be assessed [6]. As the conoid of the ookinetes does not extrude *per se*, it implies that the role of the PCRs in *Toxoplasma* and *Plasmodium* might be slightly different. The PCRs in ookinetes are always seen above the pellicular entrance by EM, suggesting that F-actin could theoretically flows continuously inside the pellicle. Interestingly, this constant F-actin flux is supported by the observations that ookinetes motility is uninterrupted and relies on a "continuous motile force" [10, 11].

Unlike in Toxoplasma, the disappearance of the PCRs upon *Pb*Pcr4 or *Pb*Pcr5 deletion could not be visualized by EM as the ookinete collar is too electron-thick (data not shown).

*Conservation of Pcr6 and Pcr1 in Apicomplexa*

Depletion of Pcr6 led to the detachment of PCRs from the conoid in *T. gondii* with a remaining point of attachment when assessed by EM. Given its conservation across the phylum, the protein plausibly fulfils a similar role in the malaria parasites.

Pcr1 was recently documented by 3D-SIM as localizing as a small apical punctum [1], while U-ExM uncovered here a clear semi-ring above the conoid. Depletion of Pcr1 did not impact on parasite survival, contrasting with the fitness conferring score assigned in the CRISPR-Cas9 screen [12]. This discrepancy could be investigated further by generation of a knockout line. Pcr1 semi-ring was always observed on the opposite side of the ICMTs anchor point, yet this protein is conserved in *Plasmodium* species that seemingly lack ICMTs. Its function at the PCRs still remains to be determined.
